# The claustrum-medial prefrontal cortex network controls attentional set-shifting

**DOI:** 10.1101/2020.10.14.339259

**Authors:** Leon Fodoulian, Olivier Gschwend, Chieko Huber, Sophie Mutel, Rodrigo F. Salazar, Roberta Leone, Jean-Rodolphe Renfer, Kazadi Ekundayo, Ivan Rodriguez, Alan Carleton

**Author notes:** These authors contributed equally to the work and are listed in alphabetical order (Leon Fodoulian, Olivier Gschwend, Chieko Huber, Sophie Mutel, Rodrigo F. Salazar). Senior authors contributed equally to the work. Correspondence should be addressed to A.C. or I.R.

## Abstract

In various mental disorders, dysfunction of the prefrontal cortex contributes to cognitive deficits. Here we studied how the claustrum (CLA), a nucleus sharing reciprocal connections with the cortex, may participate in these cognitive impairments. We show that specific ensembles of CLA and of medial prefrontal cortex (mPFC) neurons are activated during a task requiring cognitive control such as attentional set-shifting, i.e. the ability to shift attention towards newly relevant stimulus-reward associations while disengaging from irrelevant ones. CLA neurons exert a direct excitatory input on mPFC pyramidal cells, and chemogenetic inhibition of CLA neurons suppresses the formation of specific mPFC assemblies during attentional set-shifting. Furthermore, impairing the recruitment of specific CLA assemblies through opto/chemogenetic manipulations prevents attentional set-shifting. In conclusion, we propose that the CLA controls the reorganization of mPFC ensembles to enable attentional set-shifting, emphasizing a potential role of the CLA-mPFC network in attentional dysfunctions.

## INTRODUCTION

Cognitive dysfunctions, which are a hallmark of many psychiatric conditions including schizophrenia and attention-deficit hyperactivity disorder (ADHD), represent a major burden in the daily life of patients (Millan et al., 2012). It is particularly true since current medications poorly alleviate this class of symptoms (Husa et al., 2017; Swartz et al., 2008). Therefore, a better understanding of the brain circuits involved in cognitive functions and/or dysfunctions is critical to better handle psychiatric pathologies and develop new therapies. Prefrontal cortex (PFC) regions have long been recognized as a neuronal hub contributing to various cognitive functions such as planning, attentional processes and working memory (Roberts et al., 1998). Recent circuit tracing studies in rodents provided a comprehensive picture of the diverse neuronal types forming synapses on various PFC neuronal populations (Ahrlund-Richter et al., 2019), which include the claustrum (CLA) as a presynaptic partner. In this work, we evaluated the potential role played by the CLA on PFC function during cognitive tasks.

A century ago, the CLA was described as a thin sheet of gray matter located between the insular cortex and the putamen (Landau, 1919). Because of its dense reciprocal connections with the neocortex (Druga, 1968; LeVay and Sherk, 1981; Minciacchi et al., 1985; Smith et al., 2012; Wang et al., 2017), the CLA has been proposed to be involved in consciousness (Crick and Koch, 2005; Koubeissi et al., 2014), saliency detection (Remedios et al., 2010, 2014), or neural synchrony (Smythies et al., 2012, 2014). Recently, several lesions or opto/chemogenetic manipulations of claustro-insular neurons suggested a role played by the CLA in contextual fear conditioning (Kitanishi and Matsuo, 2017), spatial reversal learning and working memory (Grasby and Talk, 2013), impulsivity (Liu et al., 2019), top-down action control and visuospatial attention (Liu et al., 2019; White and Mathur, 2018), attention (Atlan et al., 2018; Goll et al., 2015; Mathur, 2014; White and Mathur, 2018; White et al., 2018), resilience to an auditory distractor (Atlan et al., 2018), as well as sleep (Luppi et al., 2017; Narikiyo et al., 2020; Norimoto et al., 2020; Renouard et al., 2015). The CLA might thus be implicated in multiple functions, though some of these suggested roles are not supported by tissue-specific CLA manipulations.

In the present study, we molecularly distinguished CLA neurons from neighboring striatal and insular cortical neurons using single cell RNA sequencing (scRNAseq) and used a Cre-driver transgenic mouse line to specifically study CLA glutamatergic projection neurons. Using conditional viral tracing, microendoscopic calcium imaging and opto/chemogenetic manipulations during cognitive tasks, we demonstrate the involvement of the CLA-mPFC network in attentional set-shifting.

## RESULTS

### Widespread axonal projections of VGLUT2^+^ CLA neurons to the cortex

To identify CLA neurons at the molecular level, we performed scRNAseq from adult wild-type brains injected in the motor cortex with an adeno-associated virus (AAV) that drives the expression of a CRE-GFP fusion protein in the nucleus of infected cells. This allowed anterograde trans-synaptic tagging (Zingg et al., 2017) of neurons in the striato-insular area (**Figure 1A**). CLA neurons formed an isolated population, which was transcriptionally distinct from neighboring subpopulations of striatal and cortical neurons (**Figures 1B, 1C** and **S1**). Consistent with recent mouse brain scRNAseq studies (Rosenberg et al., 2018; Saunders et al., 2018; Zeisel et al., 2018; Zeisel et al., 2015), a clustering analysis revealed that a single neuronal population expressed several CLA marker genes such as *Gng2, Lxn, or Oprk1* (Wang et al., 2017). The expression patterns of identified marker genes found in our clustering and genes known to be expressed in the claustro-insular area were compared between each other (**Figure 1D**). In the mouse brain, CLA neurons are located in very close apposition with other neuronal populations of the insular cortex (expressing for example *Nos1ap*, *Sdk2* or *Ctgf*) and olfactory areas such as the endopiriform nucleus (EN) and the piriform cortex (expressing for example *Nnat* and *Ctgf*; **Figure 1D**). The vesicular glutamate transporter VGLUT2 (encoded by the *Slc17a6* gene), which was previously proposed to be selectively expressed in the CLA but not in surrounding areas of adult rodent brains (Hur and Zaborszky, 2005), was one of the most selective markers of CLA neurons. Its expression enabled to differentiate the CLA from the nearby EN (**Figure 1D**, see also the discussion). Therefore, we used *Vglut2*-ires-Cre knockin mice combined with viral transfections to specifically target CLA neurons for the rest of the study.

**Figure 1.**
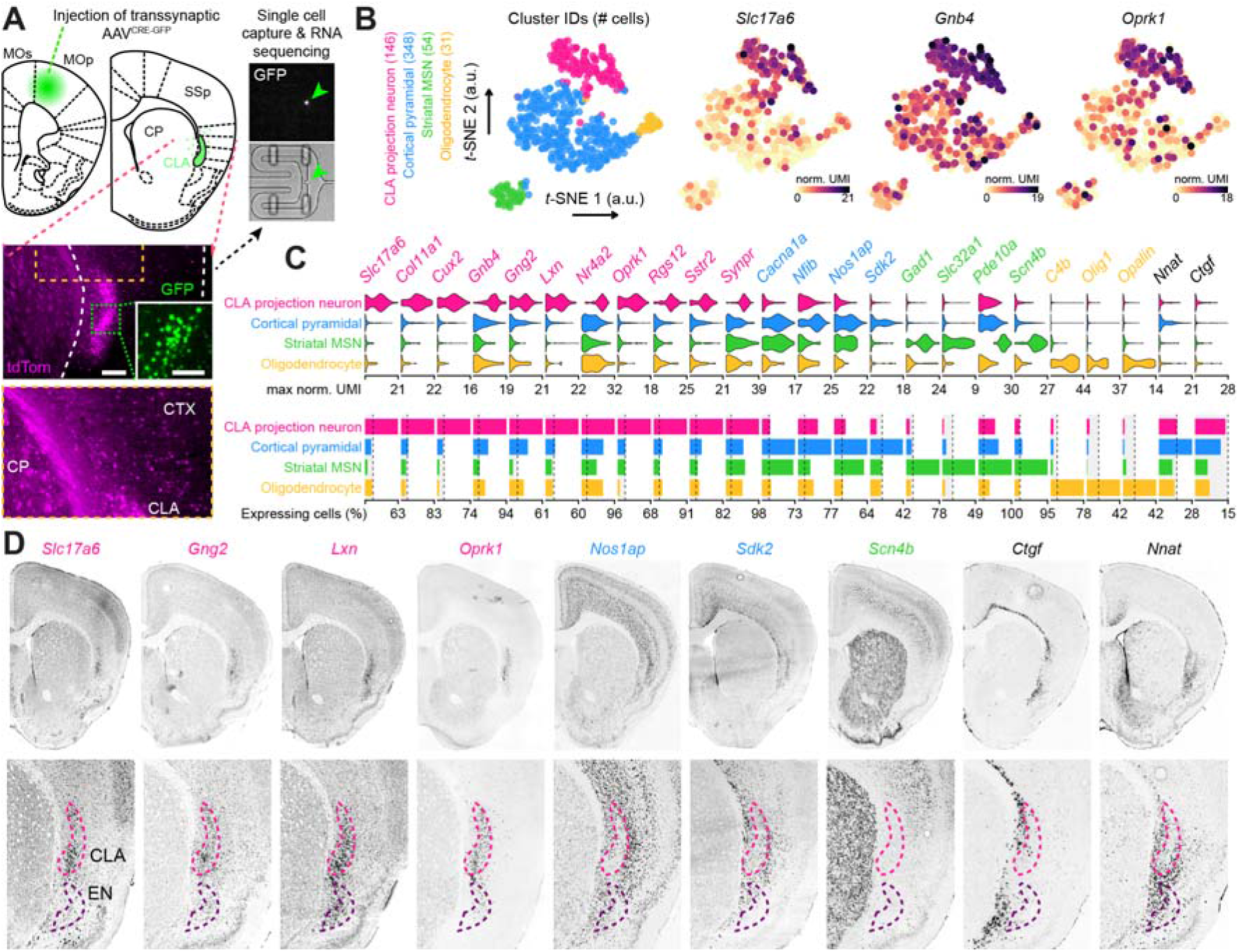
Single cell transcriptomic in the claustro-striato-insular area. (**A**) Schema of the experimental design for scRNAseq (see methods). *Bottom*: whole-cell tdTomato labelling of neurons in the claustrum-striatum-insula area and nuclear GFP-CRE localization seen at higher magnification (*inset*, scale bars: 120 & 60 μm, respectively). *Right*: Representative capture chamber from the C1 high-throughput integrated fluidic circuit. Green arrow shows a captured GFP^+^ cell. (**B**) Visualisation of cell clusters on *t*-distributed stochastic neighbour embedding (t-SNE) plot and representation of cell-specific gene expression of three CLA marker genes. (**C**) Cluster-specific marker genes. *Top*: Violin plots showing cluster-specific distribution of marker genes expression. *Bottom*: Bar plots representing the percentage of cells in each cluster expressing the corresponding gene at a minimum of 1 normalized UMI count (dashed lines are placed at 15%). Gene name colours indicate the cell type in which they are upregulated. Black colour indicates genes not upregulated in any of the clusters. (**D**) Representative in situ hybridization data of selected genes from the Allen Mouse Brain Atlas. *Top*: Coronal hemisections of the mouse brain. *Bottom*: Higher magnification view of the claustro-insular area.

We first used an anterograde viral tracing strategy to analyze the pattern of axonal projections of VGLUT2^+^ CLA neurons in the mouse brain (**Figures 2A** and **2B**). As reported with other Cre-driver transgenic lines (Atlan et al., 2017; Narikiyo et al., 2020; Wang et al., 2017), projections were widespread, targeting the large majority of cortical areas, although the density strongly varied between regions; the frontal, cingulate, temporal and insular regions presenting the highest densities (**Figures 2C** and **2D**). Overall, associative cortices exhibited higher axonal densities than sensory and motor cortices (**Figure 2D**). Furthermore, CLA axons were not homogenously distributed across the various cortical laminae (**Figure 2E**). They indeed preferentially innervated superficial rather than deep layers in most cortical areas (e.g. prelimbic and anterior cingulate cortices), though the opposite was observed for few exceptions (e.g. perirhinal and lateral entorhinal cortices; **Figures 2E-2G**).

**Figure 2.**
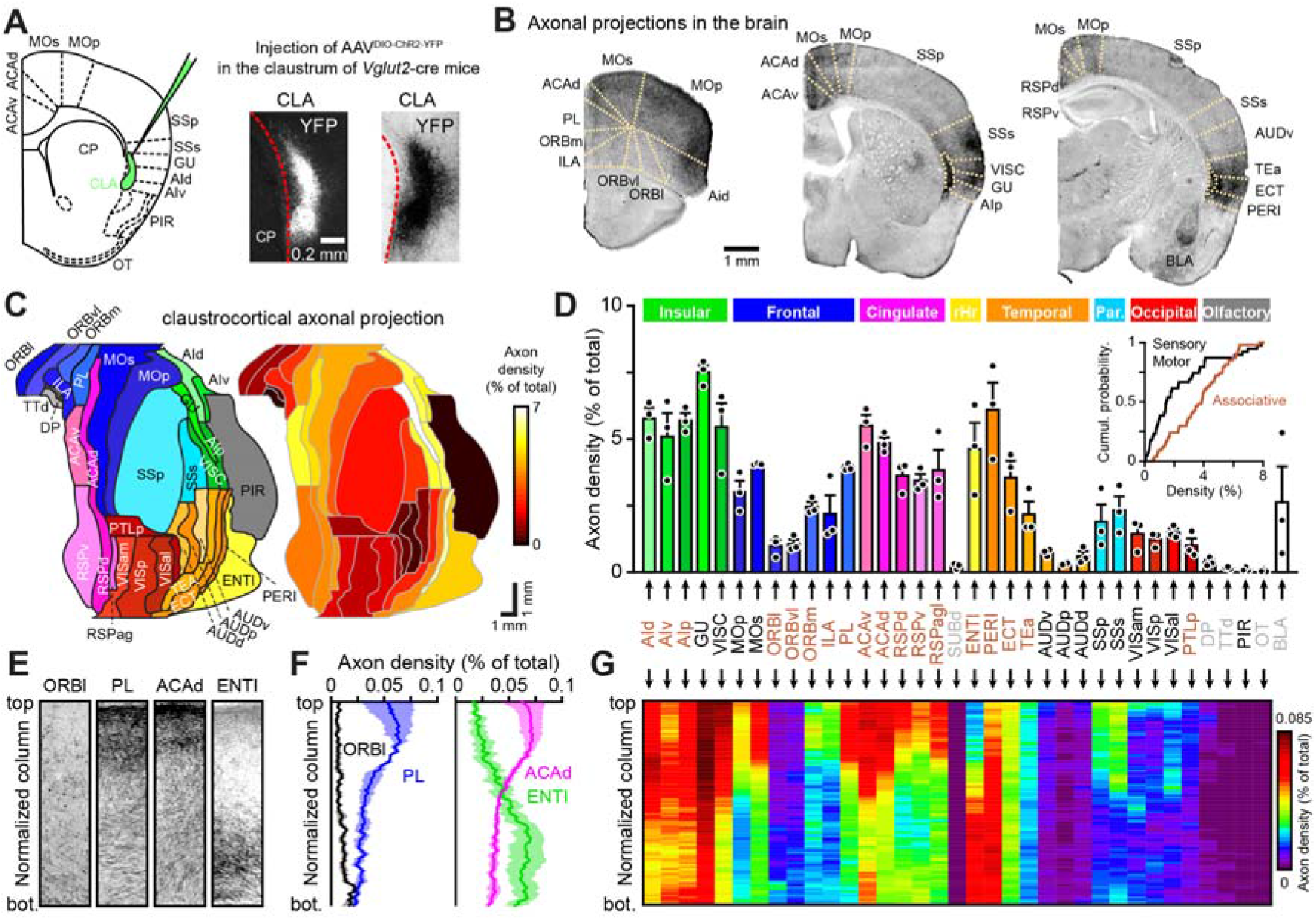
Widespread claustro-cortical glutamatergic projections in the adult mouse brain. (**A**) Schema of the experimental design for axonal tracing. *Photographs*: YFP expression in virally infected VGLUT2^+^ CLA neurons (fluorescence and DAB staining shown for two different mice, respectively). (**B**) Coronal sections of a CLA-infected brain immunostained for YFP (dark staining). (**C** and **D**) Quantification of the relative density of axonal projections to various brain areas arising from CLA-VGLUT2^+^ neurons (*n* = 3 mice). Data are presented as an unrolled cortical map (C) or as a histogram (D, black circles represents values obtained for each mouse). Densities varied between brain areas (Friedman ANOVA on all areas: *P* = 2×10D^◻7^). Inset: cumulative distribution of axon density in primary and secondary motor/sensory (black) vs. associative cortices (red; *n* = 39 and 54 area/mouse pairs, respectively, Kolmogorov-Smirnov test *P* = 0.0014). (**E**) Photographs of cortical columns (homogeneously stretched to the same length) taken from different areas showing the uneven distribution of CLA axons across layers. (**F** and **G**) Distribution of axonal projections arising from CLA-VGLUT2^+^ neurons in cortical columns of various areas (repeated measures two-way ANOVA: PL *vs.* ORBl *F*_(99, 198)_ = 6.04, *P* < 10^◻10^; ACAd *vs.* ENTl *F*_(99, 198)_ = 32.5, *P* < 10^◻10^). Top and Bot.: pial surface and bottom of the columns, respectively. Data are presented as mean ± SEM. See table S1 for brain structure abbreviations and table S2 for detailed statistics.

### Specific CLA assemblies are recruited during various cognitive states

Since frontal regions are implicated in executive functions (Birrell and Brown, 2000; Bissonette et al., 2008; Dias et al., 1996; Ng et al., 2007; Pantelis et al., 1999; Tait et al., 2009) and since they are densely innervated by CLA projection neurons in both mice (Wang et al., 2017) (**Figure 2**) and primates (Reser et al., 2016; Reser et al., 2014), we hypothesized that the CLA might be involved in cognitive flexibility and attention. To test these hypotheses, we monitored the activity of VGLUT2^+^ CLA neurons expressing the genetically encoded calcium indicator GCaMP6f using microendoscopic calcium imaging (**Figures 3A-3C**) while mice performed an attentional set-shifting task (ASST). This task challenges attentional load and cognitive flexibility through trial-and-error learning (Brown and Tait, 2016; Keeler and Robbins, 2011; Tait et al., 2014). The task is composed of a sequence of trial blocks in which sensory cues of different natures (olfactory and tactile cues are used here) have to be associated with a reward. Each of those blocks consists in a different set of stimulus-reward contingencies (**Figure 3D**), and is known to require the integrity of specific cortical areas. For example, selective lesions of the lateral orbitofrontal cortex, and of the mPFC in rodents (or their equivalent in primates), selectively impair reversal learning and attentional set-shifting, respectively (Birrell and Brown, 2000; Bissonette et al., 2008; Dias et al., 1996; McAlonan and Brown, 2003; Ng et al., 2007; Pantelis et al., 1999; Tait et al., 2009).

**Figure 3.**
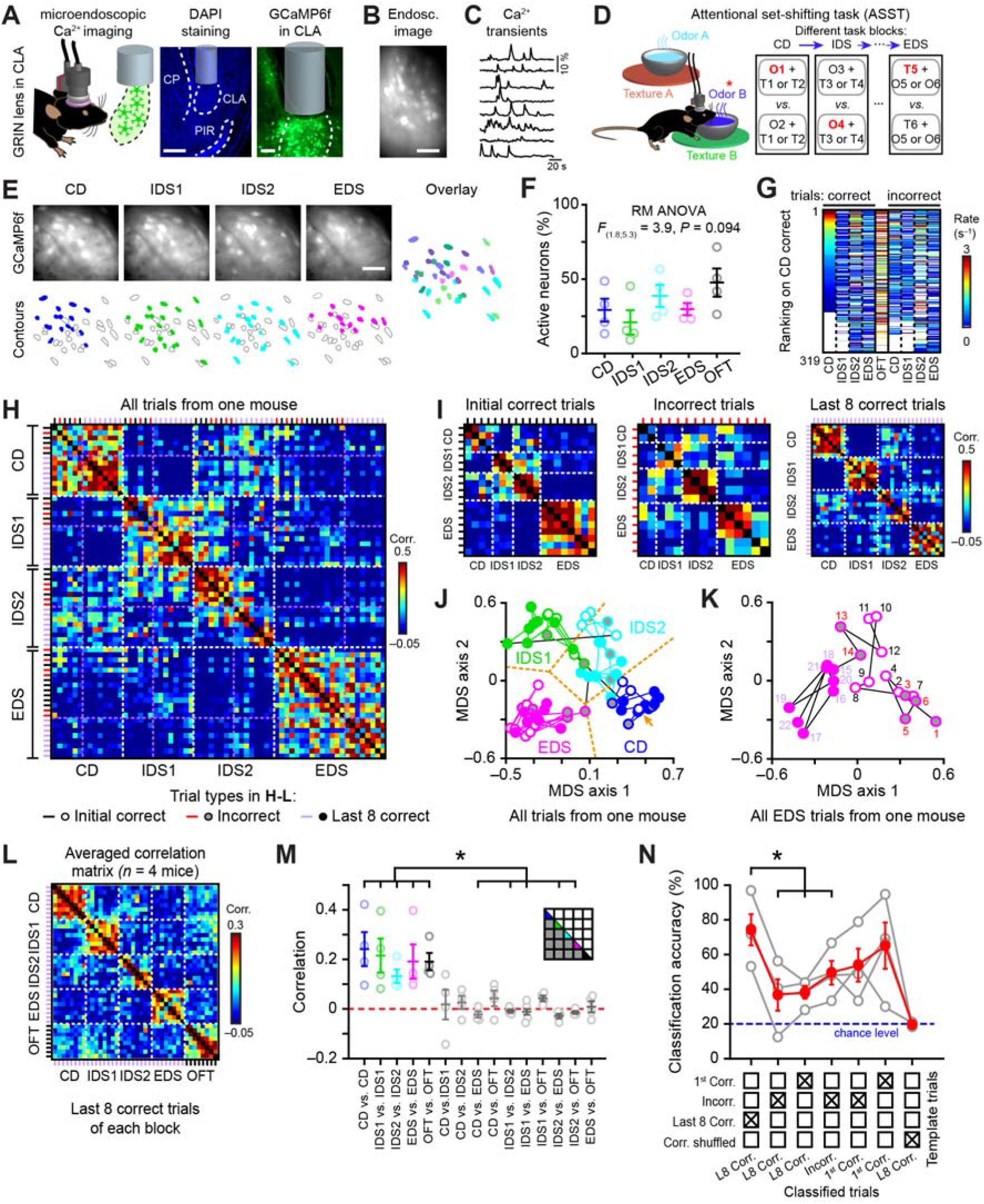
Different cognitive states recruit specific claustral cell assemblies. (**A**) Schema of the recording procedure and histological verification of GRIN lens position in the brain (CP: caudate putamen, CLA: claustrum, PIR: piriform cortex). Scale bars: 200 & 40 μm for DAPI & GCaMP, respectively. (**B**) VGLUT2^+^ CLA neurons expressing the genetically encoded calcium indicator GCaMP6f and imaged with a microendoscope (scale bar: 50 μm). (**C**) Examples of calcium transients recorded from eight CLA neurons in a freely behaving mouse. (**D**) Schema of an attentional set-shifting task procedure. Each block of the task is composed of a varying number of trials depending on animal performances. In all trials, two odorants (presented as clean odorized bedding in two pots) and two textures (presented below a pot) are randomly combined and placed in two separate compartments of a behavioral box. Only one cue (either one odor or one texture) is reinforced by a food reward hidden in the bedding. During the compound discrimination (CD) block, mice must learn that only one sensory dimension and one stimulus predicts the reward location (odor O1 in the example). After reaching a learning criterion (see methods), a new set of cues is presented and a positive transfer of the learned rule is expected to take place (formation of an attentional set); i.e. mice should make a new reward-cue association following the same relevant sensory dimension as in the previous block (intradimensional shift, IDS block). In the extradimensional shift (EDS) block, a new reward-cue association is made by reinforcing a cue within the previously irrelevant dimension. This attentional set-shifting task assesses the ability to attend to previously irrelevant information. CD and IDS1 were performed on the same day whereas IDS2 and EDS were performed on the following day. (**E**) Representative examples of CLA neuronal ensembles during the ASST. The pictures show the maximum intensity projections of imaging data collected during various blocks of the ASST. Example of specific ensembles of cells exhibiting calcium responses during each block of the task and are highlighted with specific colors in the contour plots (firing rate is above 1.5 Hz trials during each session, scale bar: 50 μm). (**F**) Percentage of active neurons during the various ASST blocks and during an open-field test (OFT, *n* = 4 mice). (**G**) Average firing rate computed in CLA neurons during an open-field test (OFT), during the last 8 correct trials (i.e. the ones defining the learning criterion) and the incorrect trials of each block of the ASST. Neurons (319 recorded in 4 mice) are ranked according to the firing rate recorded during the correct trials of the CD block, showing that specific cell assemblies are activated in different types of trials recorded during the various blocks of the ASST. (**H**) Correlation matrix showing the similarity of ensemble representations for all consecutive ASST trials imaged in one mouse (for each pair of trials, the Pearson correlation coefficient is calculated using two vectors of CLA neuron firing rate computed over the duration of the considered trials). Magenta dashed lines mark the beginning of the last 8 correct trials of each ASST block whereas white dashed lines emphasize the transitions between different ASST blocks. Incorrect, the last 8 correct and other correct trials (referred as initial correct trials) are indicated in in red, purple and black, respectively. (**I**) Correlation matrices showing the same data as in H for the different trial types. (**J** and **K**) Multidimensional scaling (MDS) analysis showing the evolution of the firing activity in the population of CLA neurons recorded in one mouse during individual trials (i.e. each circle) of either the entire ASST (J) or only the EDS block (K, numbers and colors indicate the actual trial order and the trial type, respectively). The black lines in (G) indicate transitions between blocks and the orange arrow marks the position of the beginning of the task (i.e. first CD trial). The orange dashed lines indicate the boundaries of each cluster region defined by a k-means algorithm. (**L**) Averaged correlation matrix (*n* = 4 mice) showing the similarity of ensemble representations imaged during the last 8 correct trials of each ASST block and OFT. White dashed lines delineate the different blocks. (**M**) Averaged correlation of ensemble representations during the last 8 correct trials of each ASST block and OFT (*n* = 4 mice; inset: schema of the matrix presented in (L) and corresponding color code; Friedman ANOVA *P* = 10^4^, post-hoc Dunn’s test at least \**P* < 0.05). (**N**) Classification performances as a function of the type of templates and the type of classified trials (see methods). Repeated measures ANOVA *F*_(1.82, 5.56)_ = 10.77 *P* = 0.013, post-hoc Fisher test at least \**P* < 0.05. Data are presented as mean ± SEM. See table S1 for brain structure abbreviations and table S2 for detailed statistics.

CLA neurons were active during various blocks of the ASST irrespective of trial outcome (i.e. incorrect or correct responses), and during the exploration of an open field arena (**Figures 3E, 3F**, **S2A** and **S2B**). To test the specificity of these activations, the cells were ranked according to their average firing rate, which revealed that the strength of cell activation varied in a trial type- and block-dependent manner. This result was the first indication that different CLA assemblies were recruited during the various ASST blocks and open-field exploration (**Figure 3G**). For each animal and for each trial, the activity of all neurons was further pooled in a population vector (Yamada et al., 2017). The reliability of cell ensembles was assessed by correlating pairs of vectors using Pearson correlation (**Figures 3H** and **3I**), and by applying multidimensional scaling (**Figures 3J** and **3K**). To evaluate the contamination of random firing to the patterns of activity, the parameters computed for each animal/block were compared to the distributions of shuffled data (**Figures S2C-S2H**). Pairwise trial correlations changed over the course of the task (**Figures 3H**) independently from the number of active neurons, the averaged population firing rate and the trial duration (**Figures S2I-S2K**). As seen for an example mouse, vectors were more correlated during trials of a given block than between trials of different blocks, independently of trial outcome (**Figures 3H** and **3I**). When projected in the same multidimensional scaling space (**Figure 3J**), population vectors defined a unique trajectory. It reflected smooth and short transitions between trials from the same ASST block, and abrupt changes between trials from different blocks. These shifts in neuronal ensembles occurring between blocks were corroborated across mice by higher correlations between trials of a given ASST block than between trials of different blocks (**Figures 3L, 3M**, **S2F** and **S2M**).

CLA ensemble responses did not solely reflect sensory-evoked responses associated to the cues used in a given block, since for the same cues we observed a trial-by-trial remapping until the learning of the block rule (**Figures 3K** and **S2G**). Indeed, a change of representation between the initial trials and the last correct trials (i.e. defining the acquisition of the rule and the block completion) was observed above chance for all animals (**Figure S2H**). To further support the idea that specific cell assemblies are recruited during various conditions, we used a template-matching algorithm applied on ensemble activity. We showed that single trial could be classified as part of a particular block (CD, IDS, EDS) or trial type (error, 1^st^ correct, 8 last correct) significantly better than when information was shuffled (**Figures 3N** and **S2N**). In summary, specific ensembles of CLA neurons were activated during various aspects of a cognitive task and displayed learning-associated remapping.

### mPFC and CLA neurons share similar response profiles during cognitive tasks

Given the involvement of the rodent mPFC in attentional set-shifting (Birrell and Brown, 2000; Bissonette et al., 2008; McAlonan and Brown, 2003; Ng et al., 2007; Tait et al., 2009) and considering the extensive connectivity between the CLA and the mPFC (Wang et al., 2017) (e.g. PL, ILA and ACA regions, **Figure 2**), we recorded mPFC activity during the ASST using microendoscopic calcium imaging (**Figure 4A**), and compared it to CLA activity. Neurons in mPFC were activated in all ASST blocks, including those whose performance is known to be unaffected by mPFC lesions (e.g. CD and IDS, **Figure S3A**) (Birrell and Brown, 2000; Bissonette et al., 2008; McAlonan and Brown, 2003; Ng et al., 2007; Tait et al., 2009). Similarly to CLA neurons, trial-by-trial remapping of mPFC neural activity led to specific ensembles at the end of each ASST block (**Figures 4B-4F** and **S3B-S3C**, see also **Figure S6G**). Interestingly, remapping drove more reliable assemblies during the block leading to attentional set-shifting (i.e. higher correlation in the EDS block; **Figure 4G**), which has been shown to depend on mPFC integrity (Birrell and Brown, 2000; Bissonette et al., 2008; Dias et al., 1996; McAlonan and Brown, 2003; Ng et al., 2007; Pantelis et al., 1999; Tait et al., 2009). In summary, mPFC and CLA share similar response profiles during the ASST.

**Figure 4.**
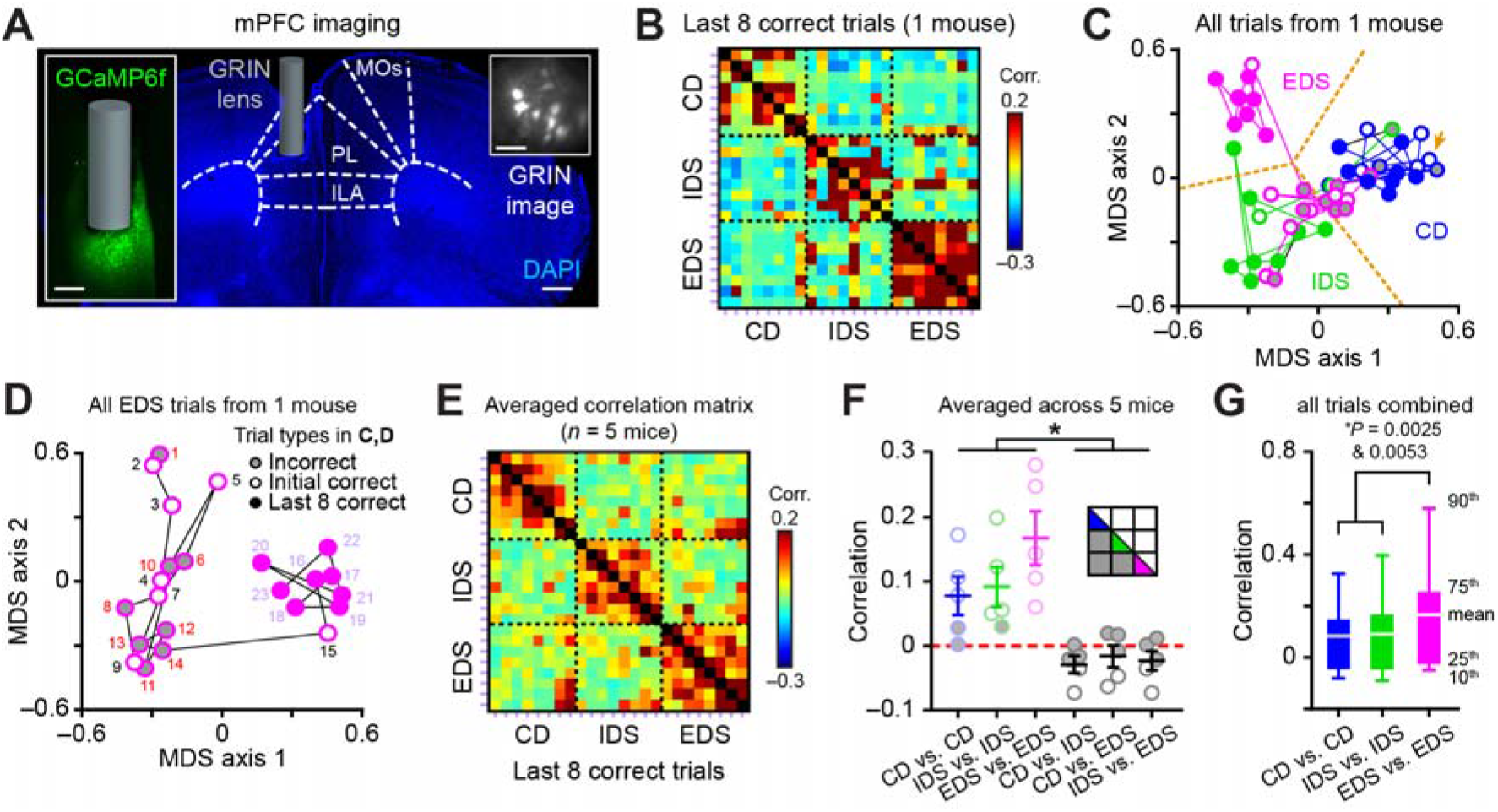
Different cognitive states recruit specific mPFC cell assemblies. (**A**) Histological verification of GRIN lens position in the mPFC (MOs: secondary motor cortex, PL: prelimbic cortex, ILA: infralimbic cortex). Photographs of mPFC neurons expressing the genetically encoded calcium indicator GCaMP6f (left inset) and imaged with a microendoscope (right inset). (**B**) Correlation matrix showing the similarity of ensemble representations imaged in one mouse during the last 8 correct trials of each ASST block. For each pair of trials, the Pearson correlation coefficient is calculated using two vectors of mPFC neuron firing rate. Black dashed lines delineate the various ASST blocks. (**C** and **D**) Multidimensional scaling (MDS) analysis showing the evolution of the firing activity in the population of mPFC neurons recorded in one mouse during individual trials (i.e. each dot) of the ASST. Note the remapping of mPFC activity during the learning of the EDS rule (D; numbers indicate the actual trial order). The black lines in C indicate transitions between blocks and the orange arrow the position of the first CD trial. The orange dashed lines indicate the boundaries of each cluster region defined by a k-means algorithm. (**E**) Averaged correlation matrix (*n* = 5 mice). (**F** and **G**) Average correlation comparing different behavioral conditions (inset: schema of the matrix presented in E and corresponding color code). Data are averaged either across mice (F, Friedman ANOVA P = 0.0011, Post-hoc Dunn’s test \**P* < 0.05, see Table S2 for detailed statistics; white symbols indicate the values which are significantly different from their respective shuffled distribution) or across data combined from all mice (G, Mann-Whitney test, *n* = 140 correlations from 5 mice for each ASST block).

### CLA glutamatergic projections increase mPFC neuron excitability

Since the CLA sends dense axonal projections to the mPFC, and since both brain structures share very similar neuronal properties during the ASST, we identified the layer-specific targets receiving synaptic connections from VGLUT2^+^ CLA neurons in the mPFC by using Channelrhodopsin-2 (ChR2) assisted circuit mapping in mPFC slices (**Figure 5A**). Stimulation of ChR2^+^/VGLUT2^+^ CLA axons evoked monosynaptic excitatory postsynaptic potentials in pyramidal neurons from various layers (**Figures 5B-5E**). We next characterized neural responses in the CLA-mPFC network *in vivo* (**Figures 5F**). We first confirmed that ChR2 stimulation successfully evoked an increase in the firing rate of optotagged CLA neurons (**Figure S4**). We explored the ability of VGLUT2^+^ CLA outputs to drive mPFC neurons (**Figures 5G, 5H, 5I**, **S5A** and **S5B**). CLA optostimulation increased the firing rate of mPFC neurons both in awake and in anesthetized mice, with a minority showing inhibition (**Figures 5J** and **5K**). Both putative pyramidal cells and interneurons, identified based on their differences in spike waveform dynamics (**Figures S5C**), displayed an increase in excitability (**Figures S5D-S5G**; see also the discussion). In summary, VGLUT2^+^ CLA neurons form functional excitatory connections that increase mPFC neuron excitability *in vivo*.

**Figure 5.**
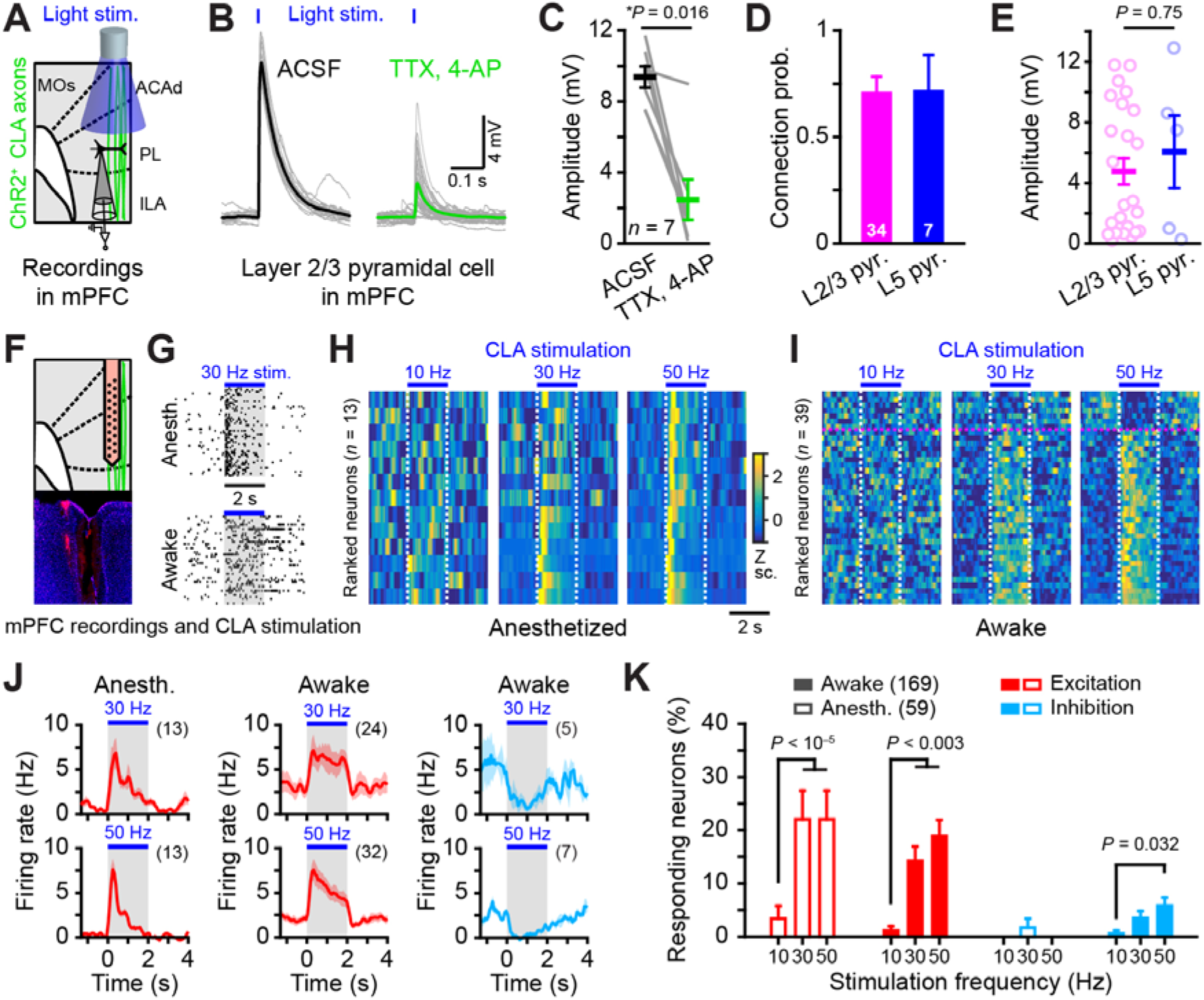
CLA glutamatergic projections increase the excitability of mPFC neurons. (**A**) Schema of the recording procedure in (B to E). mPFC slices were prepared from adult *Vglut2*-ires-cre mice previously infected in the CLA with an AAV-DIO-ChR2-YFP virus. Cortical neurons were recorded by whole cell patch clamp while CLA axons (green) were photostimulated with a blue LED (473 nm). (**B**) Example of a monosynaptic CLA connection evoked on a mPFC pyramidal cell. (**C**) Summary graph of the amplitude of light-evoked EPSPs recorded in artificial cerebrospinal fluid (ACSF) and after bath application of 1 μM TTX, 100 μM 4-AP (Wilcoxon-paired test *Z* = 2.28, *p* = 0.016). (**D**) Probability of synaptic connection made by VGLUT2^+^ CLA neurons onto mPFC neurons (recorded from 15 mice; connection probability for L2/3 *vs.* L5 pyramids: χ^2^ test *p* = 0.97). Error bars are computed assuming a binomial distribution (i.e. 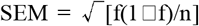, where f is the fraction of connected cells and n is the total number of recorded neurons indicated on the bars). (**E**) Summary graph of the amplitude of light-evoked EPSPs recorded in ACSF from L2/3 and L5 pyramidal cells (*n* = 24 and 5 respectively; Mann-Whitney test). (**F**) Schema of the recording procedure in (G to K). (**G**) Representative raster plots showing the response of two mPFC units to a 30 Hz CLA stimulation. (**H-J**) PSTHs for mPFC single units significantly changing their firing rate during CLA stimulations. Neurons are ranked according to their response during the 50 Hz stimulation (magenta dashed line separates units increasing from units decreasing their firing rate). (**J**) Population PSTHs averaged for mPFC single units significantly changing their firing rate during CLA stimulation (parentheses: number of responding units, colored surfaces represent the SEM). (**K**) Percentage of responsive cells to CLA stimulation at various frequencies (χ^2^ test). Neurons displaying an increase or a decrease in firing rate are presented in red or in blue, respectively. Error bars are computed assuming a binomial distribution. Data are presented as mean ± SEM. See table S1 for brain structure abbreviations and table S2 for detailed statistics.

### Impairing the activation of specific CLA assemblies prevents attentional set-shifting

We next evaluated the consequences of reducing the excitability of CLA neurons on mPFC activity. For that, we used an inhibitory designer receptor exclusively activated by designer drugs (DREADD; hM4Gi) specifically expressed in VGLUT2^+^ CLA neurons, while imaging mPFC cell assemblies during the ASST (**Figures 6** and **S6**). CLA inhibition did neither change the fraction of active mPFC neurons nor their mean population calcium rate (**Figures 6B, 6C**, **S6A-S6B**). In contrast, it modified the distribution of calcium rate of the mPFC population during the EDS block (i.e. less cells with high activity and more cells with low-intermediate activity, **Figure 6D**) and also prevented the formation of reliable cell assemblies during the EDS block (**Figures 6E, 6F**, and **S6C-S6G**). Interestingly, these mice were concomitantly unable to complete the EDS block (the other blocks being unaffected; **Figures 6G, 6H** and **S6D**), similar to what would be observed with a mPFC lesion (Birrell and Brown, 2000; Bissonette et al., 2008; Dias et al., 1996; McAlonan and Brown, 2003; Ng et al., 2007; Pantelis et al., 1999; Tait et al., 2009).

**Figure 6.**
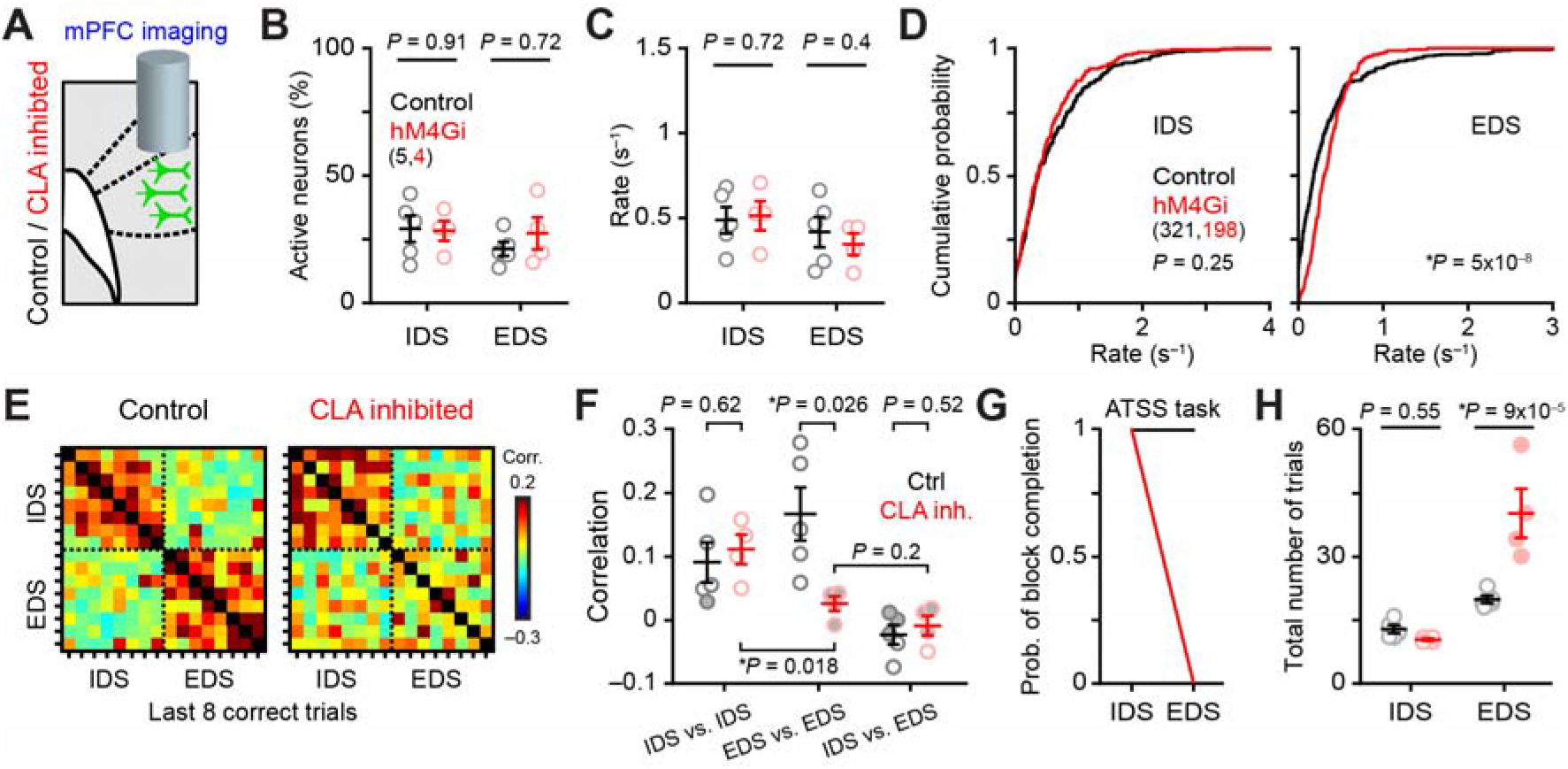
CLA glutamatergic projections promote the formation of specific mPFC cell assemblies during attentional set-shifting. (**A**) Schema of the recording procedure. Effect of CLA inhibition (by inhibitory DREADD receptor activation) on mPFC activity monitored by microendoscopic imaging during the ASST. Mice which had an AAV-Tom injected in the CLA and others which had no injection in the CLA were grouped as controls. (**B**) Percentage of active neurons (*n* = 5 and 4 mice for each group, Mann-Whitney test). (**C-D**) Effect of the CLA inhibition on mPFC neuron firing across mice (C) or across all neurons (D, Kolmogorov-Smirnov test, *n* = 371 and 198 neurons for control and CLA inhibited groups, respectively). (**E**) Correlation matrices showing the similarity of ensemble representations imaged during the last 8 correct trials of ASST blocks. Black dashed lines delineate the blocks. (**F**) Average correlation comparing different behavioral conditions (*n* = 5 and 4 mice, repeated measures two way ANOVA, [treatment x block comparison] interaction *F*_(2, 14)_ = 4.9 \**P* = 0.025; post-hoc Fisher test; white symbols indicate the values which are significantly different from their respective shuffled distribution). (**G-H**) Probability of successful block completion during the ASST in the population of tested mice (G; EDS χ^2^ test *P* = 0.0027) and number of trials performed (K; each mouse is indicated by a circle; red filled circles indicate animals unable to complete the block) during various blocks of the ASST (2 way RM ANOVA interaction [block x treatment] *F*_(1,7)_ = 17.46 *P* = 0.0042, post-hoc Fisher test). Data are presented as mean ± SEM. See table S2 for detailed statistics.

To further test the possibility that specific CLA ensembles mediated attentional set-shifting, we disrupted the formation of specific CLA assemblies by either reducing CLA neuronal activity via chemogenetic inhibition (using hM4Gi DREADD), or by increasing neuronal activity of CLA neurons using ChR2 stimulation (**Figure 7A**). None of these manipulations affected the ability to learn simple associations (e.g. SD or CD), to perform intradimensional shifts (IDS), or reversal learning (i.e. the reversal of reward contingencies for previously learned cues; e.g. CDR or IDSR; **Figure 7B**). Those results suggest that perceptual and associative abilities were not disrupted. Interestingly, an absence of effect was also reported in similar tasks after mPFC damage (Birrell and Brown, 2000; Bissonette et al., 2008; Dias et al., 1996; McAlonan and Brown, 2003; Ng et al., 2007; Pantelis et al., 1999; Tait et al., 2009). In contrast, both CLA manipulations impaired the ability of mice to perform extradimensional shifts (**Figure 7C**). This impairment was not due to a lack of motivation (ChR2 and hM4Gi groups usually performed more trials than their control cohorts; **Figure 7D**) or to alterations of locomotion or exploratory behaviors (**Figure 7E**). Overall, ChR2 and hM4Gi actuated groups were behaving at chance levels irrespective of their number of attempts (**Figures S7A** and **S7B**). Finally, a selective impairment of the EDS block was also observed when stimulating ChR2^+^ CLA terminals in the mPFC during the ASST (**Figures 7F-7H**). In conclusion, we propose that CLA ensembles are formed during specific cognitive tasks, driving the reorganization of mPFC ensembles and promoting attentional set-shifting.

**Figure 7.**
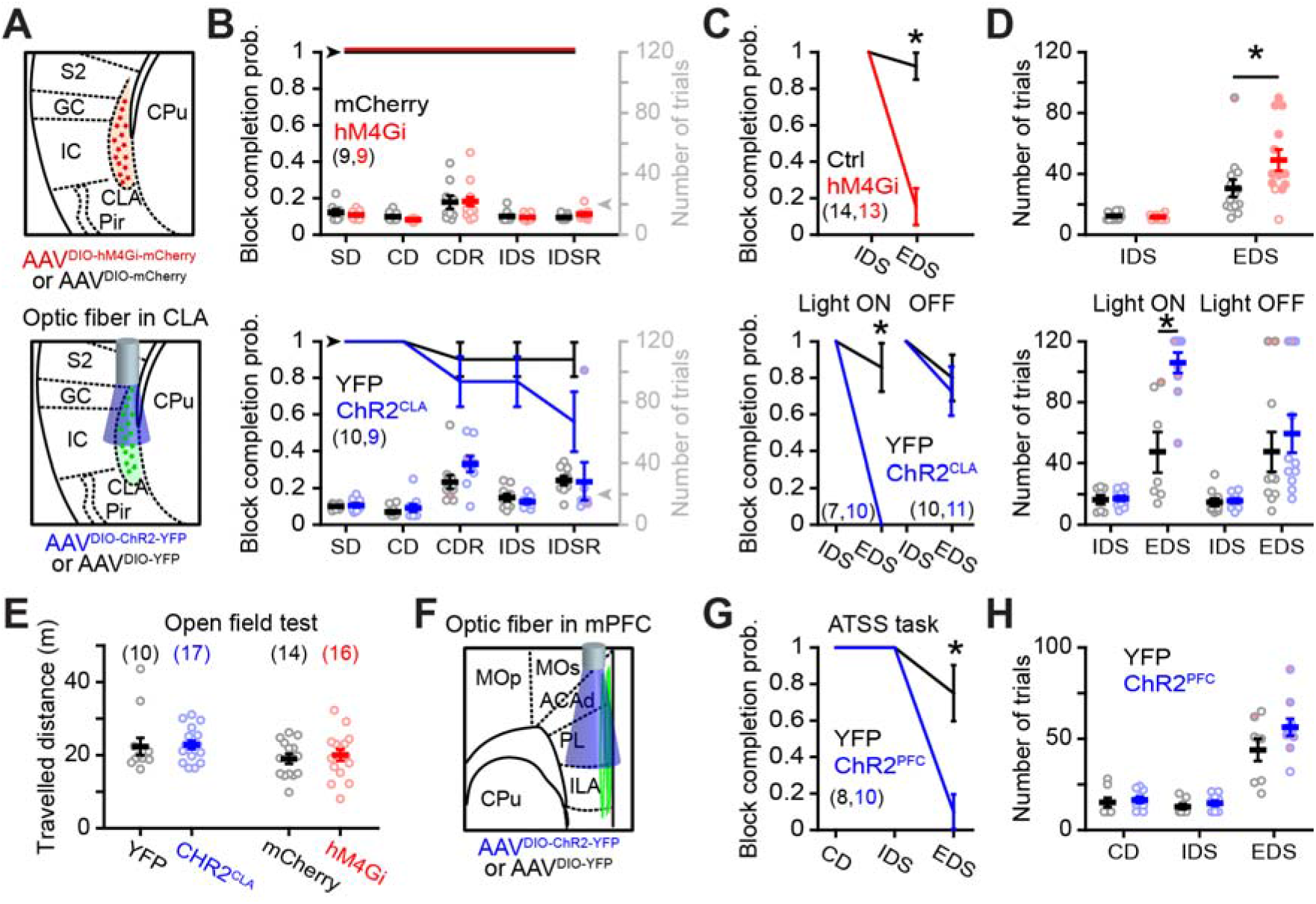
Optogenetic and chemogenetic perturbations of CLA neurons firing impair attentional but not affective shifts. (**A**) Schema of the chemogenetic (upper panels) and optogenetic (lower panels) manipulations of CLA neurons used in (B to D). (**B**) Probability of successful block completion during the ASST in the population of tested mice and number of trials performed during various blocks of the ASST (each mouse is indicated by a circle; red filled circles indicate animals that failed to complete the block). Probability of successful block completion (χ^2^ test, all *P* > 0.09). Number of trials done: mCherry *vs.* hM4Gi 2-way repeated measures (2WRM) ANOVA interaction [block x treatment] *F*_(4,64)_ = 0.27 *P* = 0.9; YFP *vs.* ChR2^CLA^ 2WRM ANOVA interaction [block x treatment] *F*_(2,34)_ = 2.4 *P* = 0.11 and *F*_(1,14)_ = 0.025 *P* = 0.88 for SD-CD-CDR and IDS-IDSR, respectively. (**C**) Probability of block completion. EDS χ^2^ test *P* = 0.000052, *P* = 0.0003, and *P* = 0.7 for mCherry *vs.* hM4Gi YFP *vs.* ChR2^CLA^ light ON, and light OFF, respectively. (**D**) Number of trials done for the blocks presented in (C). mCherry *vs.* hM4Gi: 2WRM ANOVA interaction [block x treatment] *F*_(1,25)_ = 5.26 *P* = 0.031, post-hoc Fisher test *P* = 0.93 *p* = 0.0042 for IDS and EDS, respectively (data originating from groups of mPFC imaged mice plotted in Figure S2, K and L were combined with data from non-imaged mice). YFP *vs.* ChR2^CLA^ light ON: 2WRM ANOVA interaction [block x treatment] *F*_(1,15)_ = 19.18 *P* = 0.00054, post-hoc Fisher test *P* = 0.94 *p* = 2×10^◻6^ for IDS and EDS, respectively. YFP *vs.* ChR2^CLA^ light OFF: 2WRM ANOVA interaction [block x treatment] *F*_(1,19)_ = 0.31 *P* = 0.59, post-hoc Fisher test *P* = 0.95 *p* = 0.38 for IDS and EDS, respectively. Numbers of mice analyzed are indicated in parentheses. For the probability of block completion curves, error bars are computed assuming a binomial distribution 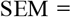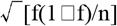, where f is the fraction of mice successfully completing a block and n is the total number of mice. Otherwise, data are presented as mean ± SEM. (**E**) Summary graph of the distance travelled during the open field test (each circle represents a mouse; Mann-Whitney test *Z* = ◻0.73 *P* = 0.47 and *Z* = ◻0.39 *P* = 0.69 for YFP *vs.* ChR2^CLA^ and mCherry *vs.* hM4Gi, respectively). (**F**) Schema of the optogenetic manipulation used in (G and H). Axon terminals of VGLUT2^+^ neurons infected with AAV2-Ef1a-DIO-hChR2-(E123/T159C)-EYFP were directly stimulated by an optic fiber implanted in the mPFC. (**G**) Probability of block completion during various blocks of the ASST. EDS χ^2^ test *p* = 0.005 (number of mice indicated in parenthesis). Error bars are computed assuming a binomial distribution 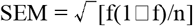, where f is the fraction of mice successfully finishing a block and n is the total number of mice. (**H**) Number of trials completed in each block (2WRM ANOVA interaction [block x treatment] *F*_(2,32)_ = 1.62 *P* = 0.21; each mouse is indicated by a circle; red filled circles indicate animals unable to complete the block; see methods).

## DISCUSSION

In this study, we demonstrate that 1) the stimulation of VGluT2^+^ CLA axons increases the excitability of mPFC pyramidal neurons both *in vitro* and *in vivo*, 2) specific CLA and mPFC assemblies are recruited during the ASST and 3) altering CLA excitability impairs the formation of specific mPFC cell assemblies and prevents attentional set-shifting (phenocopying the behavior of animals with mPFC lesions, as reported in the literature). Taken together, our data strongly suggest that the CLA-mPFC network is essential to attentional set-shifting.

### Molecular heterogeneity of neuronal populations in the claustro-insular area

Using single-cell RNA-sequencing, we molecularly distinguished claustral neurons from striatal and insular cortex neurons. The identity of each cluster was attributed according to previously identified marker genes known to characterize various neuronal populations. Except for striatal medium spiny neurons, we did not identify other GABAergic populations such as those present in the cortex or in the CLA. Whether this reflects a molecular similarity between striatal spiny neurons and other GABAergic neurons, a sampling size issue, or a capture bias in the Fluidigm chambers linked to the morphology of specific GABAergic neurons (for example, compared to CLA projection neurons and medium spiny neurons that have dense cell protrusions, GABAergic interneurons are relatively small) is unclear.

We identified 269 CLA marker genes (**Figure S1**), among which *Slc17a6* (i.e. *Vglut2*) was one of the most specific. Moreover, *Vglut2* was expressed in at least two-third of CLA projection neurons (62.3%), a proportion similar to the one characterizing several previously identified CLA marker genes such as *Gng2*, *Lxn* and *Oprk1* (**Figure 1**). These percentages are likely underestimated given the prevalence of dropout events (i.e. undetected transcripts due to failures in reverse-transcription or cDNA amplification) observed in single-cell RNA-sequencing studies (Kharchenko et al., 2014). Based on differential intrinsic electrophysiological properties, various studies proposed the existence of multiple populations of CLA projection neurons (Graf et al., 2020; Shibuya and Yamamoto, 1998; White and Mathur, 2018). In contrast, our molecular analysis identified a single population of claustral excitatory neurons. This latter result is probably not due to a sampling bias (e.g. labelling strategy or small sampling size) since a single population of CLA neurons was also reported in recent, wide scale and untargeted mouse brain single-cell RNA-sequencing studies (Saunders et al., 2018; Wang et al., 2019; Zeisel et al., 2018). One explanation for this discrepancy could be that different subpopulations of CLA projection neurons may share similar molecular profiles; the electrophysiological differences only reflecting small changes in relative channel expression, which is difficult to separate by transcriptional clustering. Alternatively, different subpopulations of CLA projection neurons could coexist, while only some of them expresses the known claustral markers. As such, the other subpopulations would be given a CLA identity essentially based on their anatomical proximity. However, the various *in situ* hybridization patterns shown in **Figure 1** in the CLA-insular area illustrate the potential problem of such an attribution. Indeed, many populations of neurons, likely belonging to different brain circuits (e.g. insular cortex, olfactory related networks such as the endopiriform nucleus or piriform cortex), are found in close apposition to each other (and even intermingled in many cases), making the identification of CLA boundaries uncertain. As an example, *Ctgf* (a known marker of layer 6b neocortical neurons) and *Nnat* are expressed by neurons found in between the external capsule and the CLA, or around the CLA. It remains unclear whether these latter cells represent different populations of neurons (e.g. layer 6b neurons projecting to the insular cortex or neurons of the EN), or whether they pertain to the CLA (note that these genes are not expressed by neurons of the scRNAseq CLA cluster). Overall, the claustral identity will have to be better defined in the future and should be ideally based on a combination of anatomical localization, molecular characteristics and connectivity. In this study, to evade this problem, we used local viral transfections in the *Vglut2*-ires-Cre mouse line to specifically target CLA neurons with a unique molecular identity.

### CLA axonal projections in the brain

Consistent with previous studies, we observed that associative cortical areas were innervated by denser CLA axonal projections compared to sensory and motor areas (Atlan et al., 2017; Wang et al., 2017; White and Mathur, 2018; Zingg et al., 2018). Most cortical layers contained CLA axons, with superficial layers usually displaying higher densities (e.g. approximatively 2 times more axons in L2/3 than in L6, see examples for mPFC areas in **Figure 2E-G**, note also that the entorhinal and perirhinal cortices exhibited an opposite distribution pattern). These results are in line with other works using other CLA marker cre mouse lines (Wang et al., 2017), but contrast with another study reporting an opposite projection pattern in layers of the mPFC areas [approximatively 2 times more axons in L6 than in L2/3, see (Jackson et al., 2018)]. This discrepancy likely results from the different approaches used to label claustro-frontal projections. Jackson and colleagues combined retrogradely transported AAV-cre injected in the mPFC of wild type mice with AAVs (driving the cre-dependent expression of a fluorophore) locally injected in the claustro-insular area. This approach is, unlike ours, not specific with regard to the genetic identity of the labelled neurons, and might tag various populations of the claustro-insular area projecting to the mPFC.

We report that VGLUT2^+^ CLA neurons form monosynaptic glutamatergic synapses on a majority of mPFC pyramidal neurons located at different cortical layers. Both in awake and anesthetized mice, this glutamatergic input increased the firing rate of PFC neurons recorded, though a few neurons were inhibited by the same stimulation. This result contrasts with previous studies reporting a long-lasting inhibition of most PFC neurons resulting from the direct recruitment of GABAergic PFC interneurons (Jackson et al., 2018; Narikiyo et al., 2020). However, this discrepancy is consistent with the unequal projection patterns described in the previous paragraph. In other words, these differences suggest the co-existence of parallel pathways projecting from the claustro-insular area to the mPFC, each one characterized by unequal layer-specific projection patterns and having opposite cortical responses. Future investigations will be required to molecularly distinguish, and further dissect these different circuits.

### CLA-mPFC network contribution to attentional set-shifting

Given the role played by the PFC in cognition and its reciprocal connectivity with the CLA, we assessed their involvement in cognitive control by recording spiking-induced calcium transients during an attentional set-shifting task. This task is highly translational between species (such as rodents, non-human primates and humans) and has been proposed to be clinically relevant to the study of various mental disorders including schizophrenia (Ceaser et al., 2008; Leeson et al., 2009; Pantelis et al., 1999) and ADHD (Halleland et al., 2012; Luna-Rodriguez et al., 2018). It is composed of subsequent blocks of trials, and switching between those blocks requires to make new cue-reward associations. Each block has been suggested to engage specific cortical areas. For example, selective lesions of the lateral orbitofrontal cortex or the mPFC selectively impair reversal learning or attentional set-shifting, respectively (Birrell and Brown, 2000; Bissonette et al., 2008; Dias et al., 1996; McAlonan and Brown, 2003; Ng et al., 2007; Pantelis et al., 1999; Tait et al., 2009). Since each task block is usually completed in a dozen trials by the subject, single neuron response properties are often difficult to assess due to the limited statistical power resulting from the low number of events. On the other hand, since attentional set-shifting is dynamic, evolving trial by trial, to average neurophysiological responses over many repetitions does not properly describe a non-stationary process. To circumvent these limitations, we used population vector methods based on single trials (Bathellier et al., 2008; Gschwend et al., 2015; Stopfer et al., 2003). These analyses revealed two fundamental observations. First, different subpopulations of CLA and mPFC neurons were active during various ASST blocks. Second, ensemble responses became more reliable over the course of each block (**Figures S2G,H** and **S5G**). The dynamic change of activity that we observe over the course of consecutive trials within a block likely results from an update of network operations based on trial history. It starts with an exploration phase, since mice are agnostic of the cue-reward association. The consequence is a sequence of random error and hits, with the latter being reinforced. During leaning, mice will develop the strategy that maximizes reward uptake and will slowly disregard the unreinforced stimulus.

Cell assemblies within the mPFC have been reported in mice performing cognitive tasks (Dejean et al., 2016; Durstewitz et al., 2010; Sirota et al., 2008), which is in line with our mPFC calcium imaging results. Interestingly, we found similar properties of cell assemblies in the CLA and in the mPFC during the attentional set-shifting task, probably reflecting the strong reciprocal connectivity between frontal regions and the CLA (**Figure 2**). The formation of mPFC and CLA assemblies do not entail their requirement to complete a particular block. In fact, we observed mPFC assemblies during CD and IDS blocks, which have been reported to be successfully completed even when this structure is lesioned (Birrell and Brown, 2000; Bissonette et al., 2008; Dias et al., 1996; McAlonan and Brown, 2003; Ng et al., 2007; Pantelis et al., 1999; Tait et al., 2009). When CLA neurons were chemogenetically inhibited, both the behavioral performance and the formation of cell assemblies in the mPFC were disrupted when a new sensory modality was reinforced (during the EDS block). However, none of these disruptions occurred in the other stages of the task. One explanation for this differential effect could be that mPFC cell assemblies were not driven by the CLA during CD and IDS, whereas they were during EDS. Interareal interactions are modulated according to cognitive demands (Gregoriou et al., 2009a, b; Salazar et al., 2004), and thereby, are likely to affect the neuronal communication (Fries, 2015) necessary to form cell assemblies spanning over different neuronal areas. Since the different blocks of the attentional set-shifting task underlie different cognitive abilities, it is likely that they drive different patterns of neuronal interactions. As a consequence, they might recruit different cells assemblies within the mPFC as well as within the neuronal areas interacting with the mPFC. Thus, the CLA influence onto the mPFC is likely to depend on the cognitive requirements, and on its related synchronized circuitry (Krimmel et al., 2019; Smythies et al., 2012, 2014; Vidyasagar and Levichkina, 2019).

Can the results of the present study be explained by a previously postulated function of the CLA? White and colleagues (2018) suggested a role of the CLA in the amplification of top-down prefrontal signals to guide action during goal-directed behaviors. In the present study, manipulations of the CLA did not alter the different stages of the task (CD, CDR, IDS and IDSR), which involve several cognitive processes such as learning cue-reward associations, inhibition of these associations, rule generalization and attentional-set formation. Therefore a general involvement of the CLA either in the control of behavioral action (White et al., 2018) or in sensory awareness (Remedios et al., 2014; Smith and Alloway, 2014) is unlikely. Finally, the CLA was proposed to play a role in resilience to sound distraction by reducing sensory responses in the primary auditory cortex (Atlan et al., 2018). Whether this observation extends to olfactory and tactile cues is unknown but it is unlikely, since our results indicate that CLA alterations do not disrupt odor and texture discriminations (see methods).

In conclusion, we propose that CLA ensembles are formed during different cognitive tasks and influence the formation of mPFC ensembles that are necessary to promote attentional set-shifting. Our findings emphasize a potential role of the CLA-mPFC network in attentional dysfunctions observed in neuropathologies such as schizophrenia (Ceaser et al., 2008; Gilmour et al., 2013; Jazbec et al., 2007; Leeson et al., 2009; Pantelis et al., 1999) and ADHD (Halleland et al., 2012; Luna-Rodriguez et al., 2018; Mueller et al., 2017).

## Acknowledgments

We thank members of A.C. and I.R. laboratories for helpful discussions and/or comments on the manuscript. We thank Chenda Kan and Véronique Jungo for expert technical assistance. We thank Christelle Cadilhac for sharing protocol for neuronal tissue dissociation. We thank the Flow Cytometry core facility, Bioimaging core facility and the iGE3 Genomics Platform at the Faculty of Medicine of the University of Geneva for expert technical assistance for the scRNAseq experiment. This research was supported by the University of Geneva, the Swiss National Science Foundation (grant numbers: 31003A_172878 to A.C. and 310030E_135910 to I.R.), the National Center of Competence in Research (NCCR) “SYNAPSY - The Synaptic Bases of Mental Diseases” financed by the Swiss National Science Foundation (grant 51NF40-185897, A.C.), the European Research Council (contract ERC-SyG-856439-CLAUSTROFUNCT to A.C. and I.R.), the Divesa Foundation (A.C. and I.R.), la fondation privée des HUG (A.C. and I.R.), the Novartis foundation for medical research (A.C. and I.R.), the Machaon foundation (O.G.) and the Claraz Foundation (L.F.).

## Author contributions

A.C. and I.R. carried out the study conceptualization and experimental design. R.F.S. established the protocol for viral infection of claustral and prefrontal neurons. O.G., S.M. and R.F.S. performed viral injections, behavioral testing, *in vivo* electrophysiological recordings and analyses. C.H. performed viral injections, microendoscopic imaging and analyses. L.F. performed viral injections, the scRNAseq experiment with some help from K.E and analyses. R.L. performed viral infections and patch clamp recordings in slices. J.-R.R. performed viral injections and immunohistochemistry. A.C. quantified the projection pattern in brain slices. A.C., I.R. and R.F.S. wrote and edited the manuscript with comments from all authors.

## STAR METHODS

### Animals

All experiments were performed on *Slc17a6*^*tm2(cre)Lowl*^/J heterozygote mice (referred as *Vglut2*-ires-cre in the text; the Jackson laboratory, strain number 016963) with the exception of the scRNAseq analysis, which used C57Bl6J WT (Charles River France) and *Gt(ROSA)26Sor*^*tm14(CAG-tdTomato)Hze*^/J mice (Madisen et al., 2010); the Jackson laboratory, strain number 007914). In *Vglut2*-ires-cre mice, the Cre recombinase is expressed in specific glutamatergic neurons, without disrupting the expression of the endogenous vesicular glutamate transporter 2 (Vong et al., 2011). Mice were housed in groups of 2–5 and maintained on a normal 12h light/dark cycle in a 24°C temperature room, with *ad libitum* access to food and water unless stated otherwise (see below specific conditions for some behavioural and imaging tests). All experiments were done during daytime. All animal protocols are in accordance with the Swiss Federal Act on Animal Protection, the Swiss Animal Protection Ordinance and were approved by the University of Geneva and Geneva state ethics committees (authorization numbers: 1007/3129/0 and GE/156/14).

### AAV injections

Mice were anesthetized with isoflurane (3–5% induction, 1–2% maintenance). The skin overlaying the skull was removed under local anesthesia using carbostesin (AstraZeneca) or Lidocaine. Mice were then head-fixed with ear-bars and a nose clamp on a stereotaxic apparatus (Stoelting). Eyes were protected from drying with artificial tears. The body temperature was monitored with a rectal probe and was maintained at ~37°C using a heating pad (FHC) during surgery. Bilateral craniotomies were performed above either the primary motor cortex (M1), the CLA, and/or the mPFC using an air-pressurized driller. Glass pipettes filled with solutions containing AAVs were inserted in one hemisphere at a time: antero-posterior (AP): 1.1 mm, medio-lateral (ML): ± 2.85 mm relative to bregma and dorso-ventral (DV): ◻2.4 mm relative to brain surface for the CLA; AP: 1.6 mm, ML: ± 0.3 mm relative to bregma and DV: ◻0.75 & ◻1.75 mm relative to brain surface for the mPFC; and AP: 1.79 mm, ML: ± 1.6 mm relative to bregma and DV: −1.0 mm relative to brain surface for M1. Different viruses (AAV2-Ef1a-DIO-hChR2-(E123/T159C)-EYFP, AAV2-Ef1a-DIO-EYFP, AAV2-hSyn-DIO-hm4D(Gi)-mCherry, AAV2-hSyn-DIO-mCherry, AAV1-Syn-Flex-GCaMP6f and AAV9-hSyn-eGFP-Cre from UNC or UPENN viral vector core facilities) were injected with regards to the different experiments (~100 nl per injection site). The pipette was left in place for a period of 3-5 min before its removal and the suturing of the skin. All infections were bilateral unless specified. Animals were then put back in their home cage to recover from the surgery. A minimum period of three weeks was allowed for viral expression before the animals underwent additional experimental procedures, except for the scRNAseq experiment where mice were euthanized 2 weeks post-infection.

### Chemogenetic manipulations

Viruses conditionally expressing either Designer Receptors Exclusively Activated by Designer Drugs (DREADDs AAV2-hSyn-DIO-hM4Di-mCherry) or control fluorescent protein (AAV2-hSyn-DIO-mCherry) were used. Gq-coupled DREADDs utilize a modified form of the human M4 muscarinic receptor (hM4Di) to induce inhibitory cellular responses in the presence of the ligand clozapine-N-oxide (CNO) (Alexander et al., 2009; Armbruster et al., 2007). Behavioral testing and/or microendoscopic calcium imaging was performed 1 hour after the first intraperitoneal (IP) injection of clozapine-N-oxide (CNO) (2 mg/kg, TOCRIS Bioscience) or after saline. When the experiments lasted more than 2 hours, CNO was repetitively injected every 2 hours.

### Single-cell capture and RNA sequencing

#### Virus injections and tissue collection

Mice were bilaterally injected in M1 with an AAV9 virus encoding a nuclear-targeted EGFP-Cre fusion protein (see above for coordinates and virus details). The serotype 9 allowed anterograde trans-synaptic tagging (Zingg et al., 2017). Pilot experiments performed with *Gt(ROSA)26Sor*^*tm14(CAG-tdTomato)Hze*^/J showed that cells were labeled in the CLA, insular cortex and striatum (native fluorescence shown in **Figure 1A**). For scRNAseq experiments, 6 week-old male C57BL/6 were used (n=8). Two weeks post-AAV injection, mice were anesthetized using isoflurane and euthanized by decapitation. Brains were immediately extracted and placed in ice-cold oxygenated artificial cerebro-spinal fluid (ACSF) containing (in mM): 124 NaCl, 3 KCl, 2 CaCl_2_, 1.3 MgSO_4_, 26 NaHCO_3_, 1.25 NaH_2_PO_4_, 10 D-glucose with an osmolarity of 300 mOsm and pH at 7.4 when oxygenated with 95% O_2_, 5% CO_2_. Coronal sections of the brain spanning the antero-posterior localization of the CLA were then cut at a thickness of 300 μm using a vibrating-blade microtome (Leica VT1000S; Leica Biosystems, Nussloch, Germany). Immediately after slicing, the CLA and the adjacent brain tissues were microdissected under an epifluorescent stereomicroscope.

#### Tissue dissociation

Tissue dissociations were performed using the Papain Dissociation System (cat #LK003150, lot #35S16330; Worthington® Biochemical Corporation, New Jersey, USA) following the manufacturer’s protocol, with slight modifications; the EBSS solutions and the medium solution with serum were adapted from (Saxena et al., 2012).

After microdissection, the tissues extracted from the 8 mice were pooled into two tubes (i.e. 2 times 4 mice) and kept in ice-cold oxygenated EBBS#1 (Saxena et al., 2012). The oxygenated EBSS#1 solution was used to reconstitute the papain and DNase I vials provided with the kit. Each of the two pools was transferred to a 50 ml Falcon tube containing 5 ml of the papain/DNase I mix, and was incubated for 20 min at 37°C in a laboratory oven (ProBlot™ Hybridization Oven; Labnet International) while rotating at a speed of 8-10 rpm. Tissues were then gently triturated with a P1000 pipette and left to decant for 1 min. Cell suspensions were carefully retrieved, passed through a 70 μm nylon cell strainer (Falcon, cat #352350) to get rid of any piece of undigested tissue, and placed in 15 ml Falcon tubes. Cell suspensions were then centrifuged (Centrifuge 5430R; Eppendorf) at 300 *g* for 5 min. Supernatants were discarded, and cell pellets were resuspended with 3ml of EBSS#2 (Saxena et al., 2012) to stop digestion. In order to remove myelin debris, two discontinuous density gradients were produced by carefully adding the 3 ml of cell suspensions on top of 5 ml of an ovomucoid-albumin protease inhibitor solution reconstituted following the manufacturer’s instructions. Tubes were centrifuged at 70 *g* for 6 min at room temperature, the supernatants discarded and the cell pellets re-suspended in 1 ml medium solution with serum. Cell suspensions were then passed through another 70 μm nylon cell strainer.

#### Fluorescence activated cell sorting (FACS)

To ensure the exclusive isolation of live nucleated cells, cell suspensions were incubated with 2 μg/ml of Hoechst 33342 (a UV fluorescent adenine-thymine binding dye; #H1399, Life Technologies™) at 37°C for 15 min. In order to exclude dead cells, 1 μM of DRAQ7™ (a far-red fluorescent DNA intercalating dye; #DR71000, BioStatus) was added to the cell suspensions before FACS sorting. GFP^+^/Hoechst^+^/DRAQ7^−^ cells were then sorted in an empty Eppendorf tube according to their forward scatter (FSC) and side scatter (SSC) properties using a Beckman Coulter MoFlo Astrios (Miami, Florida) cell sorter with a 100 μm nozzle at a pressure of 25 psi. Doublets were excluded after gating on FSC-A/FSC-H, followed by SSC-H/SSC-W. Approximately 3000 GFP^+^/Hoechst^+^/DRAQ7^−^ cells were collected from each single-cell suspension pool, each in a final volume of 8 μl.

#### Single-cell capture, cDNA library preparation and single-cell RNA sequencing

After FACS sorting, 2 μl of C1 Suspension Reagent (Fluidigm) was added to the cell suspensions, yielding mixes of 300 cells/μl. Cells were captured using the C1™ Single-Cell mRNA Seq high-throughput (HT) integrated fluidic circuit (IFC) designed for 10-17 μm cells (cat #100-5760, Fluidigm). Each 10 μl mix was loaded on one side of the chip which was processed on the C1 System (cat #100-7000, Fluidigm) following the manufacturer’s protocol. Following cell capture, the chip was imaged using an automated inverted fluorescent microscope (AxioObserver Z1, Zeiss) which allowed to locate single GFP^+^ cells in the 800 capture chambers. A custom Matlab script was used to interpolate and calculate the position and the focus in z for each of the 800 capture chambers based on those defined for the first capture chamber and about 20 capture chambers randomly distributed across the C1 chip. Cell lysis, barcoding, reverse transcription and PCR amplification were performed directly on the C1 chip using Fluidigm’s C1™ Single-Cell mRNA Seq HT Reagent Kit v2 (cat #101-3473, Fluidigm) following the manufacturer’s protocol. The cDNA content of each column on the chip (formed of 40 cells) was pooled before harvesting. Twenty cDNA libraries were generated using the Nextera XT DNA Library Preparation Kit (cat #FC-131-1024, Illumina) and Nextera XT Index Kit (cat #FC-131-1002, Illumina) following Fluidigm’s protocol. Before the final purification step, all 20 cDNA libraries were pooled. The library pool molarity and quality were assessed with the Qubit 2.0 using the Qubit™ dsDNA HS Assay Kit (cat #Q32854; Thermo Fisher Scientific) and the TapeStation using the Agilent High Sensitivity DNA Chip (cat #5067-5584; Agilent Technologies). The pool was then loaded at 7 pM on 4 lanes for clustering on a rapid paired-end Illumina Flow Cell (cat #PE-402-4002; Illumina) and sequenced on an Illumina HiSeq 2500 sequencer using the HiSeq Rapid SBS Kit v2 (cat #FC-402-4021; Illumina) chemistry. Read 1 consisted of 11 bases (6 bases for the cell barcode, and 5 bases for the unique molecular identifier), while read 2 was formed of 90 bases.

### Single-cell RNA-sequencing analysis

#### Demultiplexing

Two rounds of demultiplexing were performed on the raw Illumina data. The first round consisted in demultiplexing the Illumina indices using bcl2fastq2 Conversion Software version 2.20.0 (Illumina), which generated fastq files for each pool of 40 cells. The second round consisted in demultiplexing Fluidigm’s C1™ HT IFC indices using the C1 mRNA Seq HT Demultiplexing Script version 1.0.2 (a Perl script from Fluidigm), which generated a pair of fastq files per cell (i.e. reads 1 and 2).

#### Mapping

Digital gene expression matrices (i.e. matrices where genes are in rows, cells are in columns, and the corresponding gene counts are matrix entries) were generated following the single-cell analysis pipeline of UMI-tools version 0.5.3 (Smith et al., 2017). First, unique molecular identifier (UMI) sequences were retrieved from read 1 and appended to the read names using the *extract* function from UMI-tools. The reads in read 2 (i.e. biological reads) were then mapped to the Ensembl release 92 of the *Mus musculus* genome reference 38 (GRCm38) using STAR version 2.5.4b (Dobin et al., 2013). Only uniquely mapped reads were retained. Reads were then assigned to genes using featureCounts version 1.6.1 of the Subread package (Liao et al., 2013, 2014), and BAM files were sorted and indexed using SAMtools version 1.5 (Li et al., 2009). Deduplicated UMIs were then counted using the *count* function from UMI-tools. For each gene, UMIs that differed by 1 hamming distance were collapsed using the directional adjacency method (Smith et al., 2017). Since each UMI contained 5 base pairs, the maximum number of UMIs per gene was expected to be 1024.

#### Data filtering

Single-cell RNA sequencing analyses were performed on R version 3.4.3. Only cells from capture chambers containing a single GFP^+^ cell were used for downstream analyses, which resulted in 595 cells. Additional cell filtering criteria were applied following a recently published paper (Mayer et al., 2018). 1) We removed all cells characterized by less than 1000 expressed genes. 2) We removed all cells for which mitochondrial counts exceeded 10% of their total counts, as high mitochondrial counts indicate suffering or dead cells. 3) We removed all cells for which the total number of reads, the total number of detected genes, the total number of UMIs, and the percentage of mitochondrial counts were three median absolute deviations away from the median, after log_10_ transformation. 4) We removed all cells that showed unusually high or low number of UMIs given their number of reads, after log_10_ transformation, by fitting a loess curve (*loess* function of the stats R package, with span = 0.5 and degree = 2) where the number of UMIs is taken as response variable and the number of reads as predictor. Those cells for which the model residual was not within three median absolute deviations of the median were filtered out. 5) We also removed all cells that showed unusually high or low number of genes given their total number of UMIs, also after log_10_ transformation, following the above-mentioned criteria. No cells were removed after steps 1 and 2. About 3% of the cells were removed during steps 3-5, which left 579 cells for downstream analyses.

In addition to cell filtering, we also selected genes based on two criteria: 1) Only protein coding genes were used for the analysis. Those genes were selected before cell filtering. 2) Genes not expressed in at least 5 cells at an expression threshold of a minimum of 1 UMI count were excluded from the dataset, after cell filtering. This filtering resulted in a total of 14299 protein coding genes.

#### Selection of high dropout genes

In order to identify distinct cellular populations, we selected genes for downstream analysis based on their mean expression and dropout-rate relationship using the M3Drop R package version 3.09.00 (Andrews and Hemberg, 2018). We fitted a depth-adjusted negative binomial (DANB) model on the raw counts, which accounts for sequencing depth and UMI tagging efficiency differences between the cells, using the *NBumiFitModel* function. The top 1430 genes (accounting for 10% of the total number of genes in the dataset) that showed significantly higher dropout rates than what is expected by the model were selected for downstream analyses using the *NBumiFeatureSelectionCombinedDrop* function.

#### Data normalization and confounder regression

To account for technical variability (i.e. tissue dissociation effects and sequencing depth differences between the cells) in downstream analyses, data normalization and confounder variable regression were performed using the *RegressOutNBreg* function of the Seurat R package version 2.3.1 (Mayer et al., 2018; Satija et al., 2015). We fitted a negative binomial model with regularized theta (i.e. overdispersion parameter) on the raw counts by modeling the UMI counts for each gene as a function of the total number of reads, the total number of UMIs divided by the total number of detected genes, and the percentage of mitochondrial counts of each cell (Mayer et al., 2018). Regularized theta estimation was performed for all genes, but the negative binomial regression was applied only on the high dropout genes (see Selection of high dropout genes). The Pearson residuals of the model, which were clipped at a range of [−30, 30], were then used for dimensionality reduction and clustering.

In (**Figures 1B-1C)**, raw gene expression values were normalized by the total number of counts per cell and scaled to 10^4^ (i.e. corresponding to norm. UMI). Plotted gene expression values are actual normalized values, but the scales are in log_10_ after adding a pseudocount of 1.

#### Dimensionality reduction and unsupervised graph-based clustering

Prior to cell clustering, principal component analysis (PCA) on the expression matrix of the high dropout genes (see Selection of high dropout genes) was performed using the irlba R package version 2.3.2 (Baglama and Reichel, 2005) implemented in the *RunPCA* function of Seurat. Only the first 50 PCs were computed. To select for significant PCs that account for the highest variance in the data, we used the *JackStraw* function of Seurat. Over 1000 iterations, we randomly selected 1% of the high dropout genes (see Selection of high dropout genes), performed a gene-wise shuffling of their expression values (i.e. shuffling cell identities) and computed their corresponding PC loadings (we will refer to these as “fake” gene loadings). A p-value for each gene-PC pair was then calculated as the proportion of absolute fake gene loadings of the PC that are larger than the absolute real loading of this pair. We then used the *JackStrawPlot* function of Seurat to calculate a p-value for each PC. This function implements a proportion test (*prop.test* function of the stats R package) where, for each PC, the number of genes having a p-value below 10^−5^ is compared to what is expected by chance following a uniform distribution of p-values. Using a feedforward selection approach, we then selected all PCs that were significant until reaching a non-significant PC. This resulted in 4 significant PCs that were subsequently used for clustering and t-distributed stochastic neighbor embedding (t-SNE).

Seurat’s *FindClusters* function was used for unsupervised clustering of the cells and cell type identification. First, the 15 exact nearest neighbors were calculated for each cell (using the *nn2* function of the RANN R package version 2.5.1) and a shared nearest neighbor (SNN) graph was computed (Arya et al., 1998; Bentley, 1975; Mount and Arya, 1998). Then, cells were clustered using the standard modularity function of the smart local moving (SLM) algorithm(Waltman and van Eck, 2013) using several resolution parameters (i.e. a sequence from 0 to 2 with increments of 0.1). The clustree R package version 0.2.0 (Zappia and Oshlack, 2018) was used to visually inspect cluster stability at the various resolutions. To avoid over-clustering of the cells, a resolution of 0.4 was used for cell population identification, resulting in a total of 5 clusters.

The Barnes-Hut implementation of t-SNE was computed using the Rtsne R package version 0.13 (Maaten, 2014; Maaten and Hinton, 2008) implemented in the *RunTSNE* function of Seurat. The first 4 significant PCs were used for t-SNE calculation. The perplexity parameter was set to 30, and the theta (i.e. speed/accuracy trade-off parameter) was set to 0.5. We used t-SNE solely to visualize the cells (i.e. cluster assignment of the cells and their expression profile of selected genes) on a 2-dimensional plot.

#### Cluster validation and differential gene expression analysis

Validation of the clustering results was performed by comparing the number of differentially expressed genes separating each pair of clusters to a set of predefined criteria (see below). Pairwise gene expression analyses between the obtained clusters were performed using the *NegBinomRegDETest* function of Seurat. This function identifies differentially expressed genes using a regularized negative binomial log-likelihood ratio test, where the overdispersion parameter theta is estimated for each gene. Similar to data normalization and confounder regression (see above), we used the total number of reads, the total number of UMIs divided by the total number of detected genes, and the percentage of mitochondrial counts of each cell as confounder variables. Genes were considered as cluster specific markers if they were expressed by at least 40% of the cells from the corresponding cluster, and had a log2 fold-change above 2 and an adjusted p-value below 0.05. Cell clusters that differed by at least 20 differentially expressed genes were considered as separate clusters. If two cell clusters differed by less than 20, but more than 10, differentially expressed genes, we considered both clusters as distinct if one of the clusters was characterized by at least 5 differentially expressed genes, and the other by the remaining genes separating both clusters. This analysis resulted in the merging of two of the clusters corresponding to cortical pyramidal neurons, leaving us with 4 final clusters. The merging of these two clusters was corroborated by the *AssessNodes* function of Seurat. This function calculates the out of bag (OOB) error of each cluster split using a random forest classifier (Wright and Ziegler, 2017). Setting an OOB of 0.15 for a random forest classifier trained at each cluster split using all the expressed genes in our dataset also resulted in the merging of the two clusters.

To identify cluster specific marker genes after cluster validation, we computed the same differential expression test described above, but by comparing the cells from a given cluster to all other cells in our dataset. Cluster marker genes were identified as genes expressed by at least 20% of the cells from the corresponding cluster, and having a log2 fold-change above 1 and an adjusted p-value below 0.05.

### Immunohistochemistry and quantification

Ten weeks-old *Vglut2*-ires-cre male mice were injected with AAV2-Ef1a-DIO-hChR2-(E123/T159C)-EYFP in the CLA. Animals were anesthetized ~3 weeks later by an intraperitoneal (i.p.) injection of pentobarbital (200 mg/kg) and perfused transcardially with 20 ml of phosphate-buffered saline (PBS, 0.1 M, pH 7.3) followed by 50 ml of 4% paraformaldehyde (PFA) in 0.1 M PBS at 4 °C. The brains were then removed and left overnight in 4% PFA. After embedding the brains in 4% agarose, 40 μm coronal slices were cut with a vibratome (Leica VT S1000) and collected in PBS (0.1 M). For immunostaining, slices were rinsed in TBST (Tris-buffered saline with tween) and TBSTT (Tris- buffered saline with tween and triton). Slices were then incubated with 10% bovine serum for 1 h at 21-25 °C and then with the primary antibody rabbit anti-YFP (1:1000, Invitrogen, catalogue number A11122), overnight at 4 °C. The day after, slices were rinsed and incubated with biotinylated goat anti-rabbit IgG (1:200, Invitrogen, catalogue number B2770) for 1 h at room temperature. Slices were then processed using an avidin– biotin–peroxidase complex (ABC kit, Vector Laboratories) and reacted with the chromogen 3,3’-diaminobenzidine (DAB, Sigma-Aldrich). Slices were mounted with DePeX mounting medium (Gurr).

Detection and quantification of axonal projections were performed as previously described (Grider et al., 2006; Sathyanesan et al., 2012). In brief, images were acquired using a camera (Microbrightfield) mounted on an Olympus microscope (using a 4x objective; pixel size 1.55 μm; all images were captured with the same exposure time and camera gain). Images of a given brain slice were then stitched together using the software plugin “Stitching” (Preibisch et al., 2009) running on ImageJ (version 1.50b). The stitched tiff images were converted to 8-bit and the lookup table was inverted in order to get labeled axons appearing bright on a dark background. Axons were then automatically detected by first applying the Hessian filter of the software plugin FeatureJ (https://imagescience.org/meijering/software/featurej/; using the “Largest eigenvalue of Hessian tensor” option and setting the “Smoothing scale” factor to 0.5 (Sathyanesan et al., 2012)). The resulting Eigen images were then converted to 8-bit binary images using the default threshold parameters in ImageJ and were further used for quantification (Grider et al., 2006; Sathyanesan et al., 2012). Cortical and non-cortical areas of the mouse brain were defined according to the 2011 coronal atlas of the mouse brain available on the Allen brain institute website (http://atlas.brain-map.org/). For each cortical area, a rectangular region of interest (ROI, width of 300 pixels) was drawn from the cortical surface until the base of the cortical column using the ImageJ’s “line tool” (with exception of SUBd and BLA for which polygonal ROIs were drawn using the “surface tool”). Axon density was calculated as 1) the average pixel intensity in the entire ROI (using the “Measure” tool of ImageJ, for **Figures 2C-2D**) and 2) the average pixel intensity along the cortical column (for the distribution across layers using the “Plot Profile” tool of ImageJ, for **Figures 2F-2G**). For each animal and for each cortical area, the quantification was performed on 2-to-9 different coronal slices and the values were further averaged to obtain a single value and a single laminar profile per area. In order to accommodate for small variations of cortical column length, each laminar density distribution was spatially binned to give a 100 bins distribution. To account for potential variations in primary viral infection, all densities are reported as a percentage of the total quantified density across areas.

In **Figure 2D** inset: the following areas were combined in “the primary and secondary sensory or motor cortices” group and the associative cortices group, respectively: (AUDd, AUDp, AUDv, GU, Mop, MOs, PIR, SSp, SSs, VISC, VISal, VISam, VISp) and (ACAd, ACAv, AId, AIp, AIv, ECT, ENTl, ILA, ORBl, ORBm, ORBvl, PERI, PL, PTLp, RSPagl, RSPd, RSPv, TEa). See **Supplementary Table 1** for the abbreviations list.

### Slice electrophysiology

Experiments were performed on 12-13 weeks-old *Vglut2*-ires-cre mice of both sexes (infected with AAV2-Ef1a-DIO-hChR2-(E123/T159C)-EYFP 2-3 weeks before). Animals were anesthetized using isoflurane before decapitation. The brain was quickly extracted and 300 μm-thick coronal slices were cut with a vibratome (Leica VT S1000, Germany) in an ice-cold oxygenated cutting solution containing (in mM): 83 NaCl, 2.5 KCl, 0.5 CaCl_2_, 3.3 MgSO_4_, 26.2 NaHCO_3_, 1 NaH_2_PO_4_, 22 D-glucose and 72 sucrose. Slices were transferred to normal artificial cerebro-spinal fluid (ACSF) at 34 °C for 30 min and then stored at room temperature before the experiment. The ACSF used during recordings contained (in mM): 124 NaCl, 3 KCl, 2 CaCl_2_, 1.3 MgSO_4_, 26 NaHCO_3_, 1.25 NaH_2_PO_4_, 10 D-glucose with osmolarity of 300 mOsm and pH at 7.4 when bubbled with 95% O_2_-5% CO_2_. Individual slices were transferred to the recording chamber and perfused with oxygenated ACSF. All recordings were performed at ~36-37 °C (recording chamber and temperature controller from Luigs & Neumann, Germany).

Neurons were visualized using an IR-DIC microscope (Olympus BX51, Germany). Whole-cell recordings were achieved using borosilicate glass pipettes with a resistance of 4– 7 MΩ filled with an intracellular solution containing (in mM) 120 K-gluconate, 10 KCl, 10 HEPES, 4 ATP, 0.3 GTP, 10 phosphocreatine and 0.4% biocytin, pH 7.2-7.3 and osmolarity 270-300 mOsm. The signal was amplified using Multiclamp 700A amplifiers (Molecular Devices, USA; sampling rate was 50 kHz), high-pass filtered at 4 KHz and digitized using the PulseQ electrophysiology package running on Igor Pro (Wavemetrics, USA).

For optogenetic stimulations, 10 ms long light pulses were delivered on claustral terminals in the mPFC using a LED (473 nm, Thorlabs, Germany), which replaced the halogen light source transmission of the microscope. The power after the objective was set to ~0.8 mW/mm^2^. Monosynaptic connections were recorded in tetrodotoxin (TTX, 1 μM; Latoxan) and 4-aminopyridine (4-AP, 100 μM; Sigma Aldrich).

### *In vivo* electrophysiological recordings

#### Experimental procedure

Eight to 41 weeks-old male and female *Vglut2*-ires-cre mice previously injected with AAV2-Ef1a-DIO-hChR2-(E123/T159C)-EYFP in the CLA were used for electrophysiological recordings in either the CLA or the mPFC (mice used for mPFC recordings had also been implanted with optic fibers above the CLA). After AAV infection (see section above), small bilateral craniotomies were performed either above the CLA (AP: 1.1 mm, ML: ±2.85 mm) or above the mPFC (AP: AP: 1.6 mm, ML: ± 0.3 mm). Then, these craniotomies were filled up with a biocompatible polymer (Kwikcast, World precision instruments), which was covered with super glue to prevent its removal during the following recovery period (3-4 days). In addition, a stainless-steel head-post was placed on top of the skull and its arms were embedded in a mixture of super glue (Cyberbond) and dental cement (Lang). The head-post was adjusted so that the position of the brain in the stereotaxic frame (used for surgery) and in the head-holder/micro-manipulator device (used for recordings) was relatively similar, which enables the use of the same coordinates (relative to Bregma, which was marked for polytrode positioning) in the two set-ups. A rounded ground wire (silver) cemented atop the dura-mater in a parietal craniotomy was used as the reference.

A few days prior to the recording sessions, mice were habituated to be head-restrained as described in the following: they were placed in a plastic tube and head-fixed by screwing the head-post on a custom-made holder (Gschwend et al., 2012; Gschwend et al., 2016). Mice underwent this procedure on a daily session for different time periods (10, 30 and 60 min for the consecutive first, second and third day, respectively). On the day of the experiment, mice were head-fixed, the biocompatible polymer removed and the dura-mater torn apart under anesthesia (2% isoflurane). To record claustral activity, a 32-channel optrode (Neuronexus technology, A1×32-Poly2-OA32) was inserted (AP: 1.1 mm, ML: ± 2.85 mm) vertically in the brain at a speed of ~2-3 μm/s. When the approximate CLA depth was reached, we photo-stimulated the neurons as described in the next paragraph. The optrode was lowered until reliable spike responses to the light pulses were observed.

To record activity in the mPFC, either a 32-channel optrode (Neuronexus technology, A1×32-Poly2-OA32) or a multishank polytrode (Neuronexus technology, Buzsaki 32L) was inserted at a speed of 2 μm/s at (AP: 1.6 mm, ML: ± 0.5 mm) with a 10° angle. During mPFC recordings, light stimulations of CLA through optic fibers (see section below) were regularly applied along the track to check for neuronal responses. The polytrode was inserted in the mPFC at a maximum depth of 3 mm (from brain surface) before total removal. Light-driven neuronal responses, consisting in abrupt changes in local field potentials (LFP), were generally obtained between 1 mm to 2 mm (from brain surface). These responses were first identified during anesthesia (1.5% isoflurane) and later on recorded during an awake state (the changes in LFP are less obvious in the awake state). Recordings were first performed in anaesthetized animals and, once the anesthesia was stopped for at least 30 min (no changes in the polytrode position), the state of the animal was verified by stimulating the vibrissa. Only if the animal displayed a behavioral response did we start the recordings during the awake state.

For photo-stimulations, a 473-nm laser driver (Doric) was calibrated, in a continuous mode, to obtain power magnitudes estimated at 15 (for 3 ms pulse duration) or 25 mW (for 1 ms pulse duration) at the tip of the optic fibers (200 μm diameter). The measurements were performed using a photodiode sensor (Thorlabs) coupled to a power meter (PM100D, Thorlabs). During claustral and mPFC recordings, photostimulation was applied for 1 or 2 s (1-3 ms pulses at 10, 30 and 50 Hz) and 2 s trains (3 ms pulses at 10, 30 and 50 Hz), respectively. Using a two-sided Wilcoxon paired test with a significance level at 5%, each unit was tested for its responses to the photostimulation by comparing the firing rates (one measure per trial) during a period prior and another period (same duration) after the onset of the stimulation. Usually, 20 to 30 trials were recorded for each condition after the exclusion of trials with visually identifiable artefacts. When significant responses were identified, inhibition or excitation was characterized by a lower or a higher rate during the stimulation period, respectively.

#### Signal processing and spike sorting

Broadband signals were band-pass filtered (0.1 Hz to 9 kHz) and digitalized at 32 KHz with the Digital Lynx system (Neuralynx). To analyze single unit activity, the broadband signal was high-pass filtered at 500 Hz and spikes were detected when the signal reached a negative threshold. Two different methods were used to extract spikes. In recordings originating from the CLA, the threshold was set at 4 standard deviations (STD) and the 32 channels polytrode was considered as 8 tetrodes for waveform clustering using the freely available softwares KlustaKwik and MClust for spike sorting. In recordings originating from the mPFC, spikes were extracted using a weak (2.5) and a strong (4.0) thresholds and were sorted using the python-based package named ‘Klusta’ and ‘Klustaviewa’ (Rossant et al., 2016). Single units (SUA) were considered if 1) a minimum refractory period of 1-2 ms was present in the autocorrelogram, 2) the cluster was clearly separated from all of the surrounded clusters in at least one dimension and 3) no apparent cross-correlation with the other clusters (in ‘Klustaviewa’, this correspond to a cluster quality above 0.95 and similarity values below 0.05) and 4) the spikes inside the cluster had a stereotyped waveform. All clusters that did not satisfy these criteria and that did not consist of artifacts were grouped as one cluster of multi-units.

### Optical fibers preparation and implantation

200 μm diameter optic fibers (OFs) were inserted in ceramic ferrules (Precision Fiber Products, Inc) and glued using a temperature sensitive glue (Precision Fiber Products, Inc). The tip of the OFs exceed the ceramic ferrules by 2-4 mm and was polished using a set of sand papers. Only OFs having a power loss <30 % were used in the experiments. The loss of power was quantified as the percent reduction in power when the OF-ferrules were added to the tip of the connecting optic fibers.

Two different sets of coordinates were used for implanting the OFs above the CLA. One set (AP: 1.1 mm, ML: +/− 2.9 mm, from Bregma and DV: −1.9 mm from the cortical surface) was used for animals planned for behavioral testing. A second set (AP: 1.1 mm, ML: +/– 3.4 μm from the Bregma, DV: −2.6 mm from brain surface) included a 10° angle and was used in animals that underwent electrophysiological recordings in the mPFC. The use of the 10° angle enabled the simultaneous use of OF-ferrules and polytrodes in the vicinity.

### Behavioral experiments

Behavioral experiments were performed on 12–30 week-old *Vglut2*-ires-cre male mice. All behaviors were monitored using a digital camera (Stoelting). Movie acquisition, mouse tracking and quantification of movements were performed using the ANYMAZE software (Stoelting). Photostimulation was applied as follows: a laser driver (Doric lenses, Canada) drove a 473-nm laser, which was connected to an OF (Thorlabs, Germany) having two FC/PC ports. This OF was also connected to a light splitter rotary rod (Doric lenses, FRJ_1×2i_FC-2FC_0.22) composed of two FC/PC ports on the opposite side. Plugged into these ports, two additional OFs (200 μm; Doric lenses), each ended by a glued ceramic ferrule (PCP), were adjusted on the mouse’s ferrules via ceramic sleeves (PCP). The tip of these OFs was used to measure the light power. Before each experiment, the power was calibrated to ~15 (CLA stimulation) or 10 (mPFC stimulation) mW (constant light value) at the end of the OF (before the connection to the ceramic ferrules implanted in the mouse’s head), as measured with a photodiode sensor (Thorlabs) coupled to a power meter (PM100D, Thorlabs).

#### Open-field test

Open-field arenas consisted in 46 × 46 cm homemade Plexiglas square enclosures placed on top of a white painted wood plate. The fields were isolated from the experimenter by white curtains. A session consisted in placing a mouse in the center of the field and in tracking its movements for 10 minutes. In the case of optogenetic stimulation experiments, neurons were stimulated by alternating a sequence of light pulses (trains of 5 ms pulses at 30 Hz for 1 min) with periods without photo-stimulation (1 min). Data are reported for the entire duration of the assay as no difference could be observed between light and no light periods.

#### Attentional set-shifting task

This task can be used to evaluate different cognitive abilities (Brown and Tait, 2016; Keeler and Robbins, 2011; Tait et al., 2014). The task is divided into multiple blocks in which sensory cues of different nature have to be associated with a food reward. The number and type of blocks (see below for their description) were adapted for different cohorts in order to change the duration of the task (CD, IDS and EDS in imaging experiments and optogenetic activation of the mPFC experiments, all blocks in the other experiments). Each block of the task is composed of a varying number of trials depending on animal performances (see below). In all trials (except for the simple discrimination block, see below), two odorants (presented as clean odorized bedding in two pots) and two textures (presented below a pot) are randomly combined and randomly placed in two separate compartments of a behavioral box. Only one cue (either one odor or one texture) is reinforced by a food reward hidden in the bedding. Testing was performed in a homemade chamber made of Plexiglas and white PVC (40 × 30 × 40 cm). A 15 × 30 cm area of the chamber was separated in the middle by an opaque white PVC separation, on each side of which the digging bowls and the textures placed below were found (bowls were made of white ceramic; 3 cm height x 6 cm diameter except for imaging experiments for which the dimensions were 3 cm height x 8 cm diameter allowing proper investigation with the microendoscope mounted on the head). Between trials, the mouse was transferred to a clean housing cage to allow cleaning of the test chamber and repositioning of the digging bowls and textures for the subsequent trial.

Several types of ASST blocks were done (in the following examples, the relevant cues to attend are initially odors and then textures). In a simple discrimination block (SD), mice learn to discriminate between two odors (O1 and O2), with no textures present. In the compound discrimination block (CD), two textures (T1 and T2) are added as “distractors”, one odor cue remaining reinforced. In an intradimensional shift block (IDS), new sets of textures and odors (O3 and O4; T3 and T4) are presented and a positive transfer of the previously learned rule (i.e. odors are relevant and texture are distractors) is expected to take place (referred in the literature as the formation of an attentional set). In a reversal block (e.g. CDR and IDSR), the reward-contingency of previously learned cues is reversed (sometimes referred in the literature as an affective shift; in this example, a previously non-reinforced odor would become reinforced). In an extradimensional shift block (EDS), new sets of textures and odors are introduced (O5 and O6; T5 and T6) but this time a new reward-cue association is expected to occur by reinforcing a cue within the previously irrelevant dimension (a texture in this example).

For all experiments presented in the figures, odor cues represented the relevant information to attend in order to solve the SD, CD, IDS, and reversal blocks. Conversely, texture information had to be attended by animals to solve the EDS block. Since it is known that animals and humans are not equally using the different sensory modalities to solve problems, we started all tasks using olfactory cues as being informative as it was easier for procedural learning. Indeed, we also tested other groups of mice that started with a CD block for which textures were the relevant cues (referred to as CD_texture_, data not plotted in any figure). Though all mice successfully completed the block, they took ~4 times more trials to reach the learning criterion (see below) than mice doing the same CD block using odors as relevant cues (referred to as CD_odor_: mean ± SD: 66.1 ± 15.8 *vs.* 15.9 ± 6.1 trials, range 35-86 *vs.* 10-28, *n* = 10 *vs.* 18 mice for CD_texture_ vs. CD_odor_ block, respectively; Mann-Whitney *Z* = 4.3 *P* = 1.6×10^4^; all mice were *Vglut2*-ires-cre expressing either ChR2 or YFP in the CLA [ChR2 and YFP groups were combined here] and were stimulated with a blue laser through an OF implanted above the mPFC, see details below). Due to the important difference in the number of trials necessary to complete a block, we abandoned this task design. However, we used the mice doing the CD_texture_ block to test whether CLA manipulations could impair texture discrimination per se. Both mice expressing YFP or ChR2 in VGluT2^+^ CLA neurons (laser stimulation in both groups done through optic fibers implanted above the mPFC), successfully completed the block and needed similar number of trials to reach the learning criterion (mean ± SD: 67.3 ± 18.1 *vs.* 65.3 ± 15.9 trials, range 45-86 *vs.* 35-80, *n* = 4 *vs.* 6 mice for YFP vs. ChR2 groups, respectively; Mann-Whitney *Z* = 0.11 *P* = 0.92). The latter result demonstrates that texture discrimination was not disrupted by optogenetic stimulation of CLA inputs to the mPFC.

Various spices and herbs, bought in a grocery store, were used as odorants (cinnamon, nutmeg, ginger, black pepper, white pepper, coriander, cumin, cardamom, garlic, curry madras, dried onion, curcuma, paprika, rosemary). These odorants were added to standard clean bedding to obtain the scented beddings used during testing. The textures used were cardboard, PVC, sandpaper, soft fabric, rough fabric, latex, polystyrene, bubble wrap, Swiffer cloth, foam, plastic sheets of different roughness. Each texture was cut to obtain rectangles of similar sizes (about 10 × 7 cm). These textures were matched by color to minimize visual differences during testing.

Mice were food restricted to ~90% of their initial weight and habituated to the empty chamber for 30 minutes on three consecutive days. On the fourth day, two digging bowls filled with non-scented bedding and a reward (20 mg sucrose pellets, TestDiet) were added on each side of the separating wall to train the mice to dig for a reward. Mice were handled by the experimenter for 15 minutes per day over the four days. For imaging experiments, the habituation was continued for three days with attaching the microendoscope on the head of the mice.

During testing, the odorant bedding-filled digging bowls were placed on each side of the separating wall. During the simple discrimination block (SD), the bowls were placed directly on the surface of the field; in all subsequent blocks, the bowls were placed on a material corresponding to a specific texture. The position of the digging bowls and textures were randomly changed between trials so that the mouse would not associate an odor to a texture or the correct choice to a side of the chamber. The bowl containing the correct cue contained a 2 mg food pellet. The mouse was placed on the empty side of the chamber (opposite to the digging bowls), facing the wall, and the video-recording of the trial was started. In the case of optogenetic stimulation experiments, photostimulation (30 Hz, 5ms pulses) started at the start of the trial and ended when the mouse signaled its choice. In the case of “no light” experiments, OFs were connected to the implanted ferrules, but no photostimulation was applied. The trial ended once the mouse had signaled a choice by penetrating the surface of the bedding with its snout or paws. If a correct choice was made, the mouse was allowed to consume the reward and was then returned to the resting cage; in the case of an incorrect choice, the mouse was directly returned to the resting cage. For few trials, if no choice was made after 5 minutes, the trial was considered incorrect and the mouse was returned to the resting cage and allowed to rest (see further details below).

In order for a block of the ASST to be considered successful, the mouse had to reach an 80% correct choice rate over 10 consecutive trials over which the last six trials had to be correct choices. Once these criteria were reached, the mouse moved on to the next block of the ASST task. Mice were allowed to perform a maximum of 120 trials for the ChR2 group (to detect potential slower learning abilities) and a maximum of 90 trials in case of CNO injection (to avoid potential clearance of the CNO drug). However, while many “failing” mice reached the maximum number of trials, some performed less trials because they gave up and stopped sampling. After allowing them to rest, the test was resumed. If the mice kept being uninterested by the task, we considered it a failure of the block. We verified that no matter the number of trials performed, the “failing” mice did not show any improvement and were behaving at chance levels (see **Figure S7B**). In some cases, the various blocks were spread over 2 to 3 days, as different mice took a different amount of time to complete all of the blocks.

For all ASST blocks, trial-and-error learning is occurring until a block rule is eventually acquired. We distinguished between three types of trials (“incorrect”, “initial correct”, and “last 8 correct” trials). When a mouse starts a block, it doesn’t know which cues will be rewarded and therefore will decide to dig in one of the bowls by chance/uncontrolled cues. Therefore, at the beginning of the block, the outcome of the first few trials should be at chance. Progressively, the mouse should update its strategy and improve the cue-reward-prediction. We assumed that a rule is acquired when 80% correct trials over the last 10 consecutive trials and at least 6 consecutive correct responses was reached, referred to as the learning criterion. For the data analysis, we have distinguished the “last 8 correct trials” from the incorrect trials and from all the other correct trials referred as “initial correct trials”. Note that the latter type of trials combines correct responses resulting from random choice with correct responses following a change of strategy leading to an accurate reward prediction.

The incorrect and initial correct trials are intermingled as they represent the trials during which mice were progressively learning the cue-reward associations (see for example the temporal sequence of all the types of trials done by one mouse during the ASST in the correlation matrix presented in **Figure 3H** and in the MDS projections in **Figures 3J-3K**). In control animals, learning a cue-reward association requires from a few to a dozen of trials; the number varying with the block type and the task history (EDS and reversals blocks requiring more trials than CD and IDS blocks, **Figure 7**). On average, ~20-30% of the trials were incorrect and ~20-30% are “initial correct”.

### Microendoscopic calcium imaging

#### GRIN lens implantation

Animals were anesthetized with an intraperitoneal injection of a mixture containing midazolam (8 mg/kg), medetomidine (0.6 mg/kg) and fentanyl (0.02 mg/kg). Dexamethasone (2 mg/kg) was intramuscularly injected to prevent brain swelling. Carbostesin was subcutaneously injected prior to any incision, and the eyes were protected from drying with artificial tears. The body temperature was monitored with a rectal probe and was maintained at ~37 °C using a heating pad (FHC) during surgery. The skin overlying the skull was removed and a craniotomy was performed over either the CLA or the mPFC. A 0.5 mm diameter GRIN snap-in imaging cannula (Model L for CLA and Model D for mPFC, Doric lenses Inc., Canada) was slowly lowered in the brain by ~2.45-2.55 mm for the CLA and by ~1.75-1.85 mm for the mPFC from the brain surface. The GRIN cannula was then secured to the skull using cyanoacrylic glue (Cyberbond) and dental cement. A custom-made head post was also firmly attached to the skull with cyanoacrylic glue and dental cement. Anesthesia was reversed by a subcutaneous injection of a mixture containing flumazenil (1 mg/kg), naloxone (0.12 mg/kg) and atipamezole (75 mg/kg), and carprofen (5 mg/kg) was intraperitoneally injected for analgesia. All mice were allowed to recover for at least three weeks after surgery and were then habituated to a snap-in fluorescence microscope body (OSFM model L, Doric lenses Inc., Canada) attachment and in vivo imaging for > 5 days.

#### Image acquisition and image processing

Imaging acquisition was done through a snap-in fluorescence microscope body (OSFM model L, Doric lenses Inc., Canada) mounted on the implanted GRIN cannula. The CE:YAG fiber light source power (465 nm output, Doric Lenses Inc.) was tuned to less than 2 mW. The field of view corresponding to 350 μm x 350 μm was imaged at a spatial resolution of 650 pixels x 650 pixels and at a frame rate of 6.67 Hz. ImageJ (US National Institutes of Health), Doric Neuroscience Studio (Doric Lenses Inc.) and customized MATLAB (MathWorks, Inc.) routines were used for image preprocessing as follows. For CLA imaging, CD and IDS1 ASST blocks were performed for a given mouse on the same day whereas IDS2 and EDS blocks were performed on the following day. For mPFC imaging, all blocks (CD, IDS, EDS) were done on the same day.

Images from different trials of a given a day were concatenated using the Stack Sorter plugin of ImageJ. Image frames were corrected for infocal (XY) plane brain motion using “Doric neuroscience studio” alignment system and smoothed by Kalman stack filter on ImageJ. The corrected stacks were inspected for out-of-focal plane with the following algorithm: correlation coefficients of fluorescent intensities from all pixels were calculated between a given frame and a reference image (average intensity projection of the most stable stack of the day) using CorrelationJ plugin on ImageJ. If the correlation coefficient of a frame was smaller than 0.9 then the frame was replaced by the average of the previous and the next images.

Relative changes in fluorescence of registered images were computed by *F*(*t*)/*F*_*0*_ = [*F*(*t*) – *F*_*0*_]/*F*_*0*_, where *F*_*0*_ is the mean image of the entire day stack. To identify neurons and to detect Ca^2+^ transients, we used a cell-sorting algorithm (Mukamel et al., 2009; Ziv et al., 2013) running on MATLAB. It applies principal and independent component analyses (PCA/ICA) and is coupled with a segmentation routine optimized for reducing the number of false positives. Ca^2+^ transients were identified by searching each trace for local maxima that had a peak amplitude higher than two standard deviations from the trace’s baseline (i.e. periods without calcium events).

All cells were mapped from one day to another by assembling their spatial filters onto a single image. The maximum intensity projection of all stacks of day one was used as a template, to which the stacks of other days were aligned via an image alignment using TurboReg (Thevenaz et al., 1998) and Image Stabilizer (K. Li, “The image stabilizer plugin for ImageJ,” http://www.cs.cmu.edu/~kangli/code/Image_Stabilizer.html, February, 2008) on ImageJ. Candidate cells across sessions that might represent the same neuron were semi-automatically identified by customized MATLAB routines with the following algorithm: if the distance between centroids of superimposed neurons was > 10 pixels (≈ 6μm), a candidate set of cells was considered to be separated neurons.

#### Behavior annotation

At the beginning of each trial, the behavior images were synchronized with the Ca^2+^ images by an external TTL signal from ANYMAZE interface (Stoelting). Each ASST trial was recorded from the beginning of the trial until the mouse got rewarded or returned to the waiting compartment. Mouse behavioral activities were analyzed manually frame-by-frame. All behavioral activities during ASST were annotated, including odor-sampling, texture-sampling, obtaining a reward, and other activities irrelevant to the assay. Criterial for these annotated behaviors are listed below: (1) The label *odor-sampling* starts when the body or head of the mouse touch the part of the bowl to climb the bowl and ends when either the mouse obtains its reward or the head of the mouse leaves the bowl in case of absence of reward. (2) The label *texture sampling* starts when the head of the mouse enters the texture area until the front forelimbs of the mouse leave the texture area. (3) The label *having reward* starts just after the mouse nose-pokes and then benefits from the reward (including the whole reward process). (4) Other irrelevant activities included several behaviors, such as self-grooming, immobility and subtle-movements.

#### Data analysis

The firing rate of each cell was defined by the sum of the Ca^2+^ transient amplitudes per bin size and divided by a duration of the bin size. The bin size for the ASST was the duration of each trial. The duration of each trial of ASST was defined from the beginning of the trial until just before the mouse got rewarded (duration of the consumption of the reward was excluded from the analysis). The exception was **Figures S2L-S2N** for which the activity was analyzed exclusively when animals sampled odors or textures.

Equation (1) shows Pearson’s correlation coefficient ***r***, where ***x*** and ***y*** are variables, and ***n*** is a number of the pairs. The matrix of Pearson’s correlation coefficient *r* was computed from a population vector of firing rate per trial. The average of all pairwise correlations of the last 8 correct trials within bock were quantified for each mouse (**Figures 3M, 3F**, **6F**, **S2F**, **S2M**, **6F** and **S6C**).

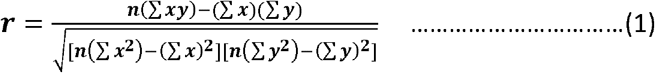

The multidimensional scaling (MDS) is one of several multivariate techniques that aim to reveal the structure of the data set by plotting points in certain dimensions. As shown in equation (2), the MDS was computed from a dissimilarity matrix ***D(r)*** obtained by the previously computed correlation matrix ***C(r**)* in this study.

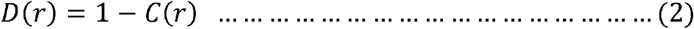

Since the MDS is similar to the PCA but the major difference is that it is based on distance among point, while the PCA is based on angles among vectors. Therefore, the MDS values varies between −1 to 1 due to the normalization within the MDS procedure. Considering that the two largest positive eigenvalues were sufficient to reproduce the dissimilarity matrix, the reconstructed trial location was plotted as a map, and *k*-means classification was applied to the map. The number of clusters was followed by the number of categories annotated type of strategy obtained by the animal strategy schema scoring and the number of blocks (k = 4 from CD, IDS1, IDS2 and EDS for **Figure 3J**, and k = 3 from CD, IDS and EDS for **Figure 4C**). The boundary was obtained by defining a fine 0.02 point distance grid on a field and using the *k*-means algorithm again to compute the distance from each centroid obtained by the *k*-means algorithm on the ASST data set to points on the grid.

The Euclidean distance between the center of all L8 trials and the center of all IC and E trials in the MDS space was computed. The first distance axis of the MDS was used as the x-axis, and the second distance axis of the MDS was used as the y-axis.

To verify whether specific cell assemblies occurred by chance and reflect the natural variance, all cells were shuffled within each trial. The shuffling was performed for 4000 iterations, and the population vector of firing rate was re-constructed at each iteration. The correlation matrix was computed based on the re-constructed population vectors. The original data and the distribution of 4000 iterations were used to find the false discovery rate, the *q* value of the false discovery rate was 5 %.

Since the activation of different cell assemblies was observed between trial types in the CLA and in the mPFC, a template-matching algorithm (Bathellier et al., 2008) was applied to test whether these cell assemblies were sufficiently reliable to classify single trials. The classification was performed based on a population vector of firing rate. The classifier was a Person’s correlation. A template population vector of firing rate was constructed from a given block of ASST (the CD, IDS1, IDS2, EDS and/or OFT). A reference population vector was constructed from trial-average traces of each block, and the trial-to-be-classified was excluded from averaging. A set of correlation coefficients were calculated between the template vectors and each candidate trial for every block. The highest correlation coefficient indicated the block to which the trial was assigned to. The accuracy/percentage of successful classification was calculated as the number of successfully classified trials divided by the total number of trials per mouse. For comparison, the classifications were repeated 1000 times on the shuffled data (cells order was shuffled within each trial and the correlation matrix was computed again).

The number of activated neurons were counted if there was at least one calcium event recorded during one trial. The percentage of neurons activated per trial was computed by the number of activated neurons divided by the total number of neurons detected during the whole ASST (**Figures 3F** and **6B**).

To evaluate whether the activation of specific cell assemblies during the different types of trials of the ASST was independent of other parameters, a regression analysis was performed between the correlation values and the duration of each trial, the number of active neurons and the calcium event rate.

### Histological verification

Except for the single cell RNA sequencing experiment, all mice were perfused transcardially with 20 mL phosphate-buffered saline (PBS) followed by 50 mL 4% paraformaldehyde (PFA) in PBS. Brains were dissected, post-fixed in 4% PFA for 24 hours, and placed in PBS. Consecutive 40 μm (for immunostaining) or 80 μm (for visual inspection of the fluorophore) coronal sections sliced with a vibrating blade microtome (LEICA VT1000S, BIO SYSTEMS) were collected onto Thermo Scientific slides, washed 3 × 3 min with PBS, sometimes stained with Hoechst (1:5000, Invitrogen, catalog number H3570) for 15 min and washed 3 × 3 min again in PBS. Slides were coverslipped with Vecta shield (Reactolab S.A.). Samples were imaged using a polarizing microscope (BX51, Olympus) using a 4X objective (UPlanFl 4X/0.13 FN 26.5, Olympus).

Usually, the fluorophore was localized within the CLA and formed a ‘peanut’ shape in both hemispheres. Occasionally, only one hemisphere (n = 15) showed some fluorescence (only one mouse had no native fluorescence observable) and the data originating from these infections were not excluded from the analyses because no obvious differences in behavior was noticed. Only rarely, some neurons located in the deep layers of the cortex in the close vicinity of the CLA were stained (n = 4). We never observed staining within the striatum.

### Statistics

All statistical analyses were performed using either R, Matlab, OriginPro 8 or Prism 7. In this study, we used ANOVA, the Wilcoxon signed-rank test, and the Mann-Whitney U test. All tests were two-sided. Unless stated otherwise, the error bars represent standard error of the mean. Data collection and animal assignment to the various experimental groups were randomized.

## SUPPLEMENTARY FIGURES

**Figure S1.**
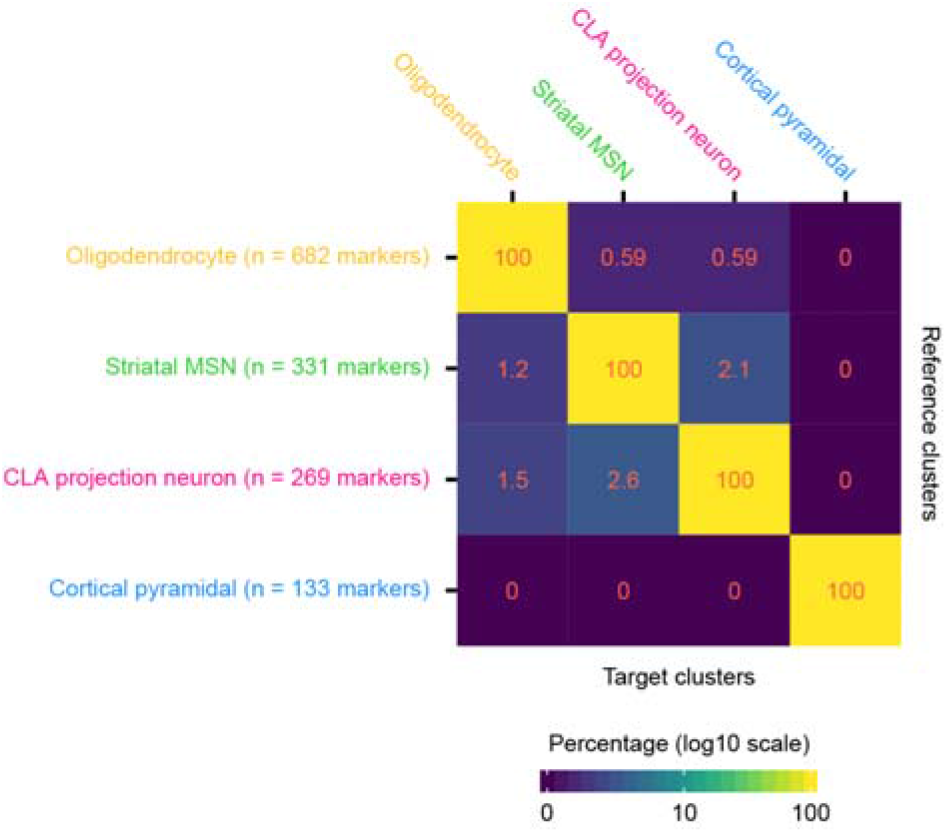
Specificity of cluster-specific marker genes, Related to Figure 1. Heatmap representation of the specificity of the identified cluster-specific marker genes. Percentages correspond to the overlap between the marker genes of reference clusters with those of each target cluster.

**Figure S2.**
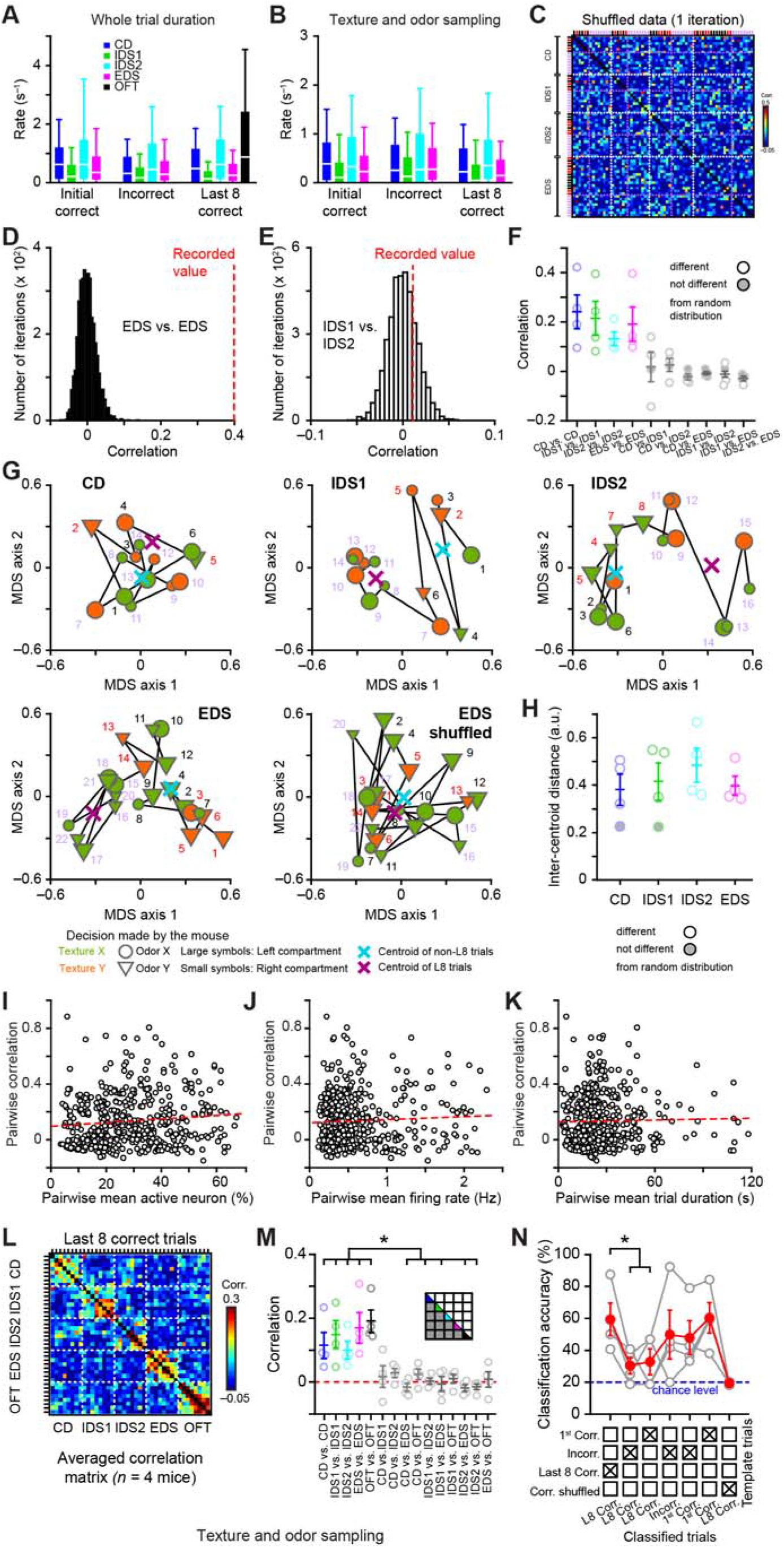
Specific claustral cell assemblies are activated during the ASST, Related to Figure 3. (**A**) and **B**) Distribution of the mean firing rate computed, either during the entire trial duration (A) or during the texture and odor sampling (B), for all CLA neurons (319 cells from 4 mice) and for different trial types (2-way RM ANOVA, interaction [ASST block x trial type] *F*_(1,318)_ = 3.83 *P* = 0.052 and *F*_(1, 318)_ = 1.38 *P* = 0.24, for A and B, respectively). (**C**) Example of a correlation matrix computed for all consecutive ASST trials imaged in one mouse (same as in Figure 3H) after one iteration of shuffled neuron identity (independently done in the ensemble for each trial). Magenta dashed lines mark the beginning of the last 8 correct trials of each ASST block whereas white dashed lines emphasize the transitions between different ASST blocks. Incorrect, the last 8 correct and initial correct trials are indicated in red, purple and black, respectively. (**D** and **E**) Two examples of mean correlation values recorded during different ASST blocks (see panel F and Figure 3H for details) and compared to their respective shuffled distribution (4000 iterations). The mean correlation value recorded in D (but not in E) is significantly different from the shuffled distribution (q < 0.05, i.e. 5% significant level with a 5% false discovery rate). (**F**) Averaged correlation of ensemble representations during the last 8 correct trials of each ASST block (each dot represents the mean correlation value for an animal, same data as in Figure 3M). White symbols indicate the values which are significantly different from their respective shuffled distribution (q < 0.05, based on analyses as in D and E). (**G**) Multidimensional scaling (MDS) analyses showing the evolution of the firing activity in the population of CLA neurons recorded in one mouse during individual trials (i.e. each symbol) of various ASST blocks. The distance between the centroids of the last 8 trials and all the other trials gives an idea of the evolution of the ensemble activity over time and can be compared to distributions of shuffled data for statistical significance (same logic as in D-F, see quantification in H). Numbers and colors (same code as in C) indicate the temporal trial sequence and each trial outcome, respectively. Each trial symbol indicates the type and location of the cues associated to the digging decision made by the mouse. (**H**) Quantification of the inter-centroid distance (as defined in G, arbitrary units) for different mouse and behavioral blocks (RM ANOVA *F*_(3,9)_ = 0.81 *P* = 0.53). White symbols indicate the values which are significantly different from their respective shuffled data, thus showing a significant change of ensemble representation over time in a given ASST block. (**I** to **K**) Ensemble correlation computed for each pair of ASST trials (from 4 mice) plotted as a function of the mean percentage of active neurons, the mean population firing rate or the mean trial duration (no significant regression). (**L**) Averaged correlation matrix showing the similarity of ensemble representations imaged during the last 8 correct trials of each ASST block (during the texture and odor sampling) and OFT (*n* = 4 mice). White dashed lines delineate the various blocks. (**M**) Average correlation comparing different behavioral conditions during the texture and odor sampling (inset: schema of the matrix presented in **g** and corresponding color code). Friedman ANOVA *P* = 3.4×10^4^, post-hoc Dunn’s test at least \**P* < 0.05. (**N**) Classification performances as a function of the type of templates and the type of classified trials during the texture and odor sampling. Repeated measures ANOVA *F*_(2.14, 6.41)_ = 5.24 *P* = 0.044, post-hoc Fisher test at least \**P* < 0.05. Data are presented as box plots in B and C (10, 25, 75, 90^th^ percentiles, median) and as mean ± SEM in (F, H, M and N).

**Figure S3.**
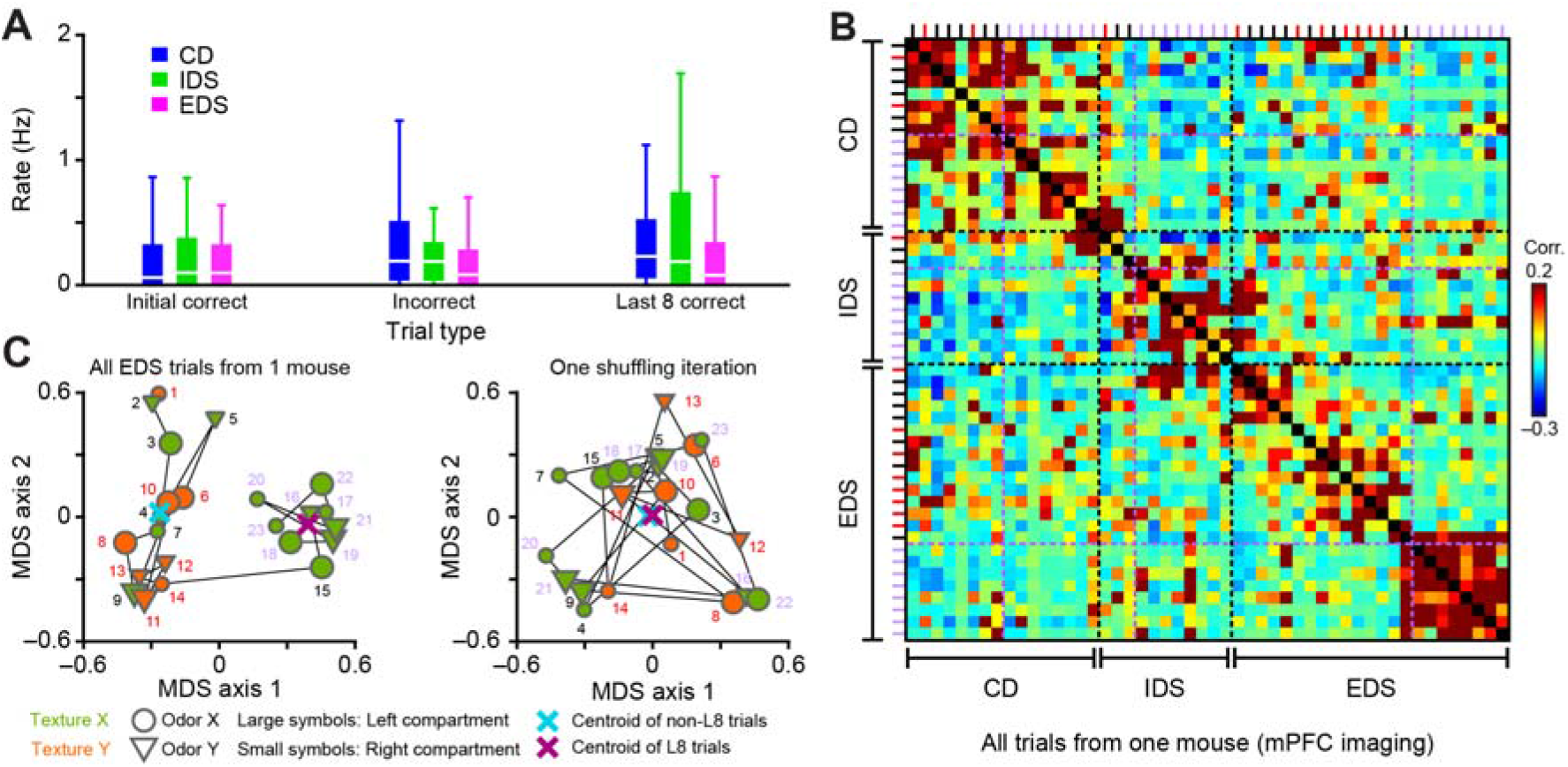
Specific mPFC cell assemblies are activated during the ASST, Related to Figure 4. (**A**) Distribution of the mean firing rate during the entire trial duration for all mPFC neurons and for different trial types (*n* = 298 neurons from 4 mice, repeated measures two way ANOVA interaction [block x trial type] *F*_(3.28, 973.6)_ = 2.17 *P* = 0.084; box plot: 10^th^, 25^th^, 75^th^, 90^th^ percentiles, median). (**B**) Correlation matrix showing the similarity of ensemble representations for all consecutive ASST trials imaged in one mouse (for each pair of trials, the Pearson correlation coefficient is calculated using two vectors of mPFC neuron firing rate computed over the duration of the considered trials). Magenta dashed lines mark the beginning of the last 8 correct trials of each ASST block whereas black dashed lines emphasize the transitions between different ASST blocks. Incorrect, the last 8 correct and other correct trials (referred to as initial correct trials) are indicated in in red, purple and black, respectively. (**C**) Multidimensional scaling (MDS) analyses showing the evolution of the firing activity in the population of mPFC neurons recorded in one mouse during individual trials (i.e. each symbol) of the EDS block (actual data and one shuffling iteration). The distance between the centroids of the last 8 trials and all the other trials gives an idea of the evolution of the ensemble activity over time and can be compared to distributions of shuffled data for statistical significance (see also quantification in Figure S5G). Numbers and colors (same code as in C) indicate the temporal trial sequence and each trial outcome, respectively. Each trial symbol indicates the type and location of the cues associated to the digging decision made by the mouse.

**Figure S4.**
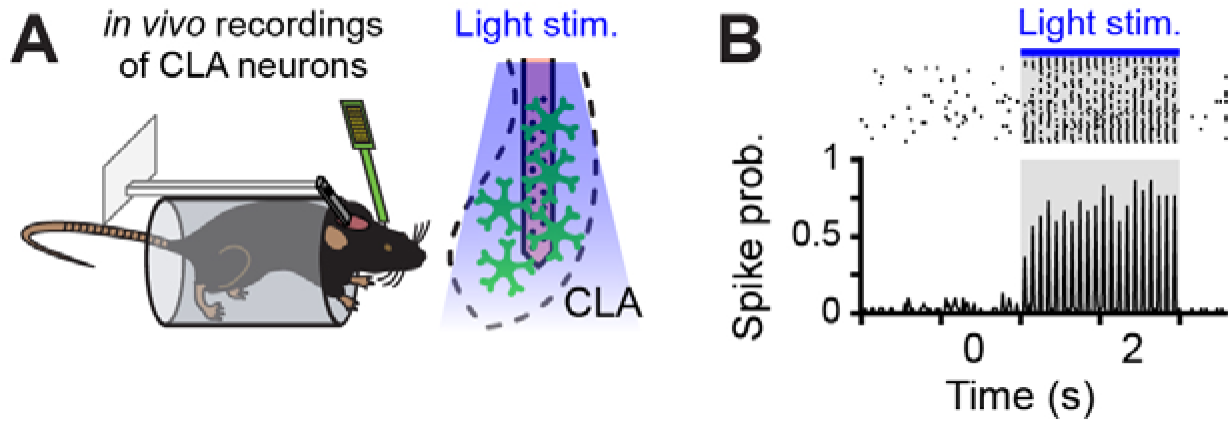
Recordings of VGLUT2^+^ CLA neurons in awake head-restrained mice, Related to Figure 5. (**A**) Schema of the recording procedure in (B). (**B**) Raster plot and peristimulus time histogram (PSTH) showing the response of a ChR2^+^ CLA unit in response to a 10 Hz light stimulation (gray boxes).

**Figure S5.**
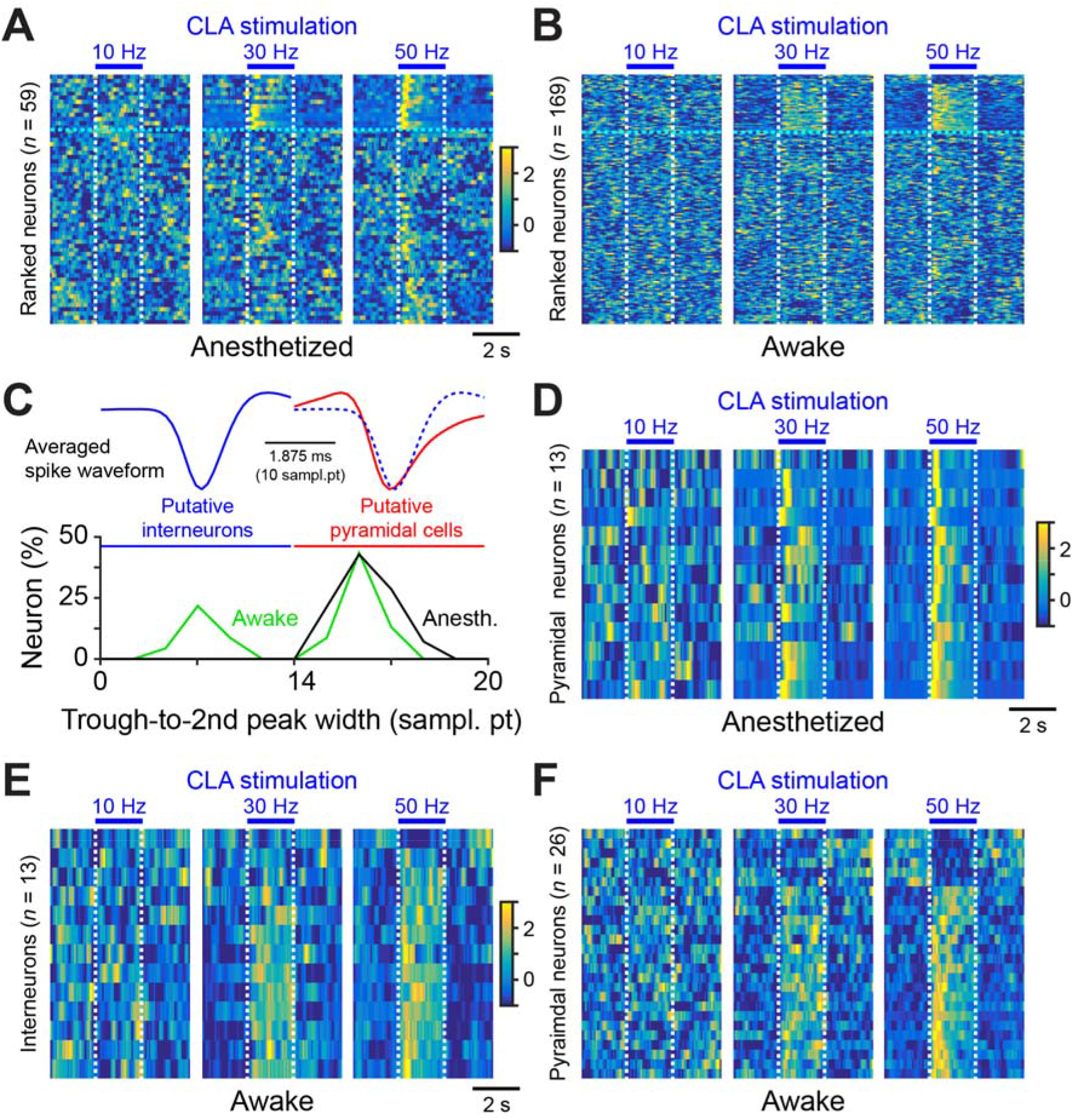
CLA glutamatergic projections increase the excitability of mPFC neurons, Related to Figure 5. (**A** and **B**) PSTHs for all recorded mPFC single units during various CLA stimulation protocols. Neurons are ranked according to their response during the 50 Hz stimulation (cyan dashed lines separate units significantly changing their firing during CLA stimulation, i.e. same neurons plotted in Figure 5H,I). (**C**) Quantification of the action potential waveform in the population of PFC units significantly changing their firing rate during CLA stimulation segregates putative pyramidal cells from putative fast spiking interneurons. (**D** to **H**) PSTHs for putative pyramidal cell and interneurons significantly changing their firing rate during CLA stimulations.

**Figure S6.**
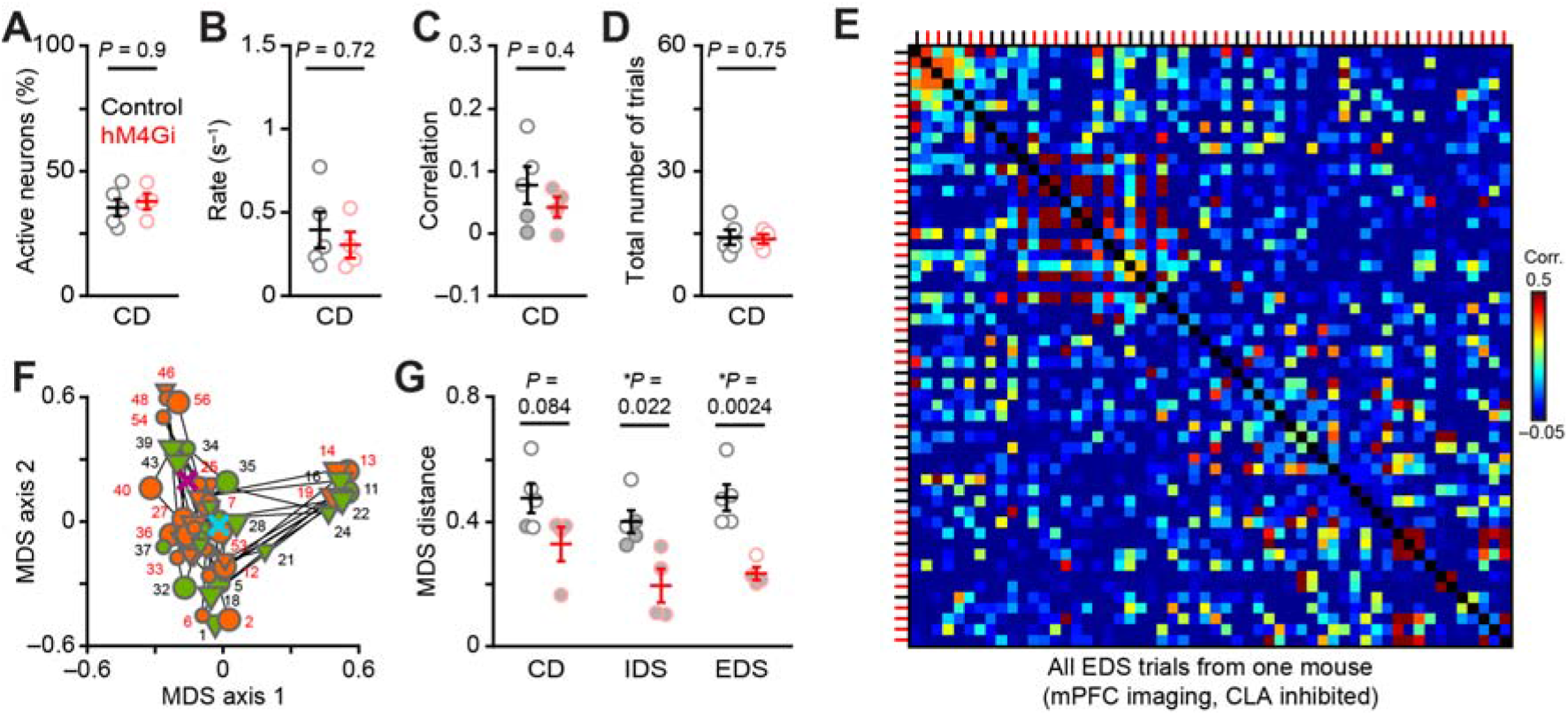
CLA glutamatergic projections promote the formation of specific mPFC cell assemblies during attentional set-shifting, Related to Figure 6. (**A** to **D**) Consequence of CLA-specific chemogenetic inhibition on mPFC neuron activity and behavior during the CD block. (**E** and **F**) Correlation matrix (E) and MDS plot (F) showing the evolution of the firing activity in the population of mPFC neurons following CLA-specific chemogenetic inhibition recorded in one mouse during individual trials of the EDS block (the mouse specifically failed to reach the learning criterion in this block, see also Figures 6 and 7). Incorrect and correct trials are indicated in red and black, respectively. Each trial symbol indicates the type and location of the cues associated to the digging decision made by the mouse. (**G**) Quantification of the inter-centroid distance (between the last 8 correct trials and all other trials) for different mouse and behavioral blocks (2 way RM ANOVA, treatment effect *F*_(1,7)_ = 164.3 *P* = 4×10^6^, Fisher post-hoc tests are indicated on the figure). White symbols indicate the values which are significantly different from their respective shuffled data. Data are presented as mean ± SEM. See table S2 for detailed statistics.

**Figure S7.**
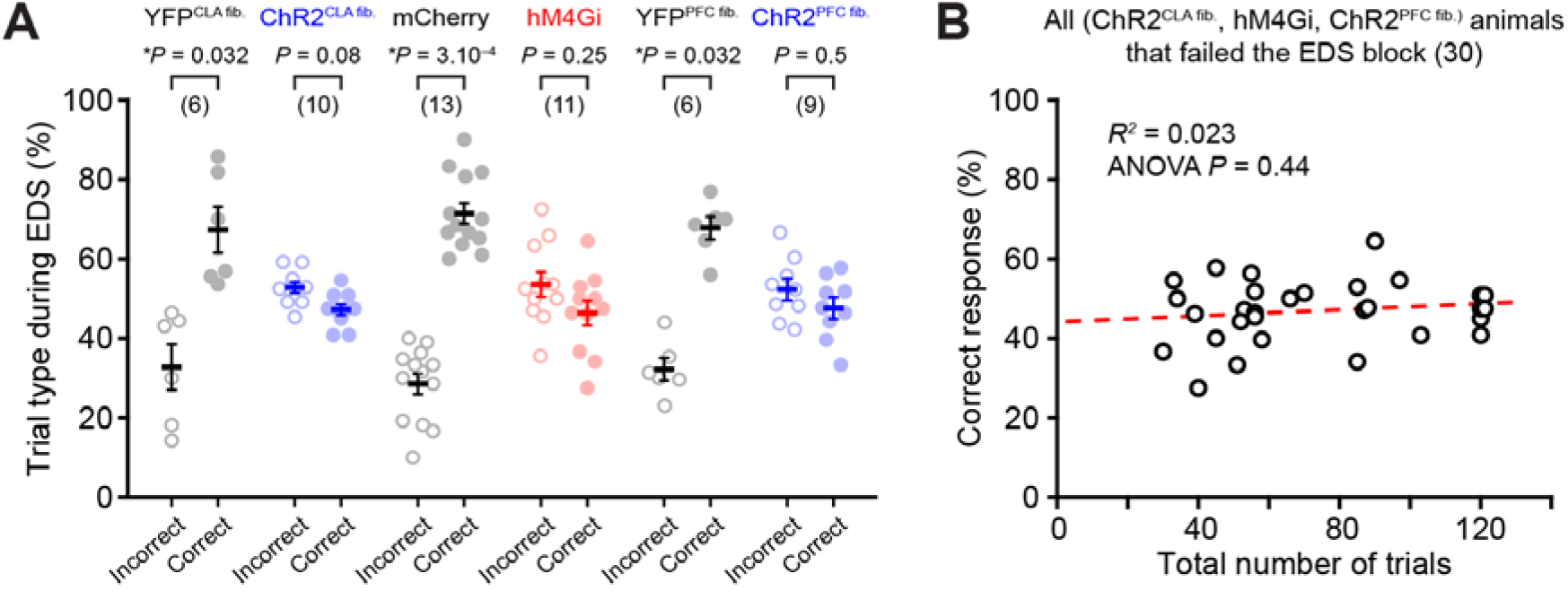
Perturbations of CLA neuron firing impair attentional set shifting, Related to Figure 7. (**A**) Percentage of trial types for control mice successfully completing the EDS block (YFP^CLA^, YFP^PFC^ and mcherry mice) and for opto/chemogenetic manipulated mice failing the EDS block (ChR2^optic fiber in CLA^, hM4Gi and ChR2 ^optic fiber in mPFC^; Wilcoxon test). We only compared control mice completing the EDS block to opto/chemogenetic manipulated mice failing the EDS block to test whether the opto/chemogenetic manipulated mice would show a learning trend, which was not the case. (**B**) Relationship between the total number of trials done and the percentage of correct trials done for all opto/chemogenetic manipulated mice that failed to complete the EDS block (combined for all experiments). Note that few data points were slightly moved to make them visible (around 120 trials). Number of animals are indicated in parentheses. Data are presented as mean ± SEM. See table S2 for detailed statistics.

## SUPPLEMENTARY TABLES

**Supplementary table 1.**
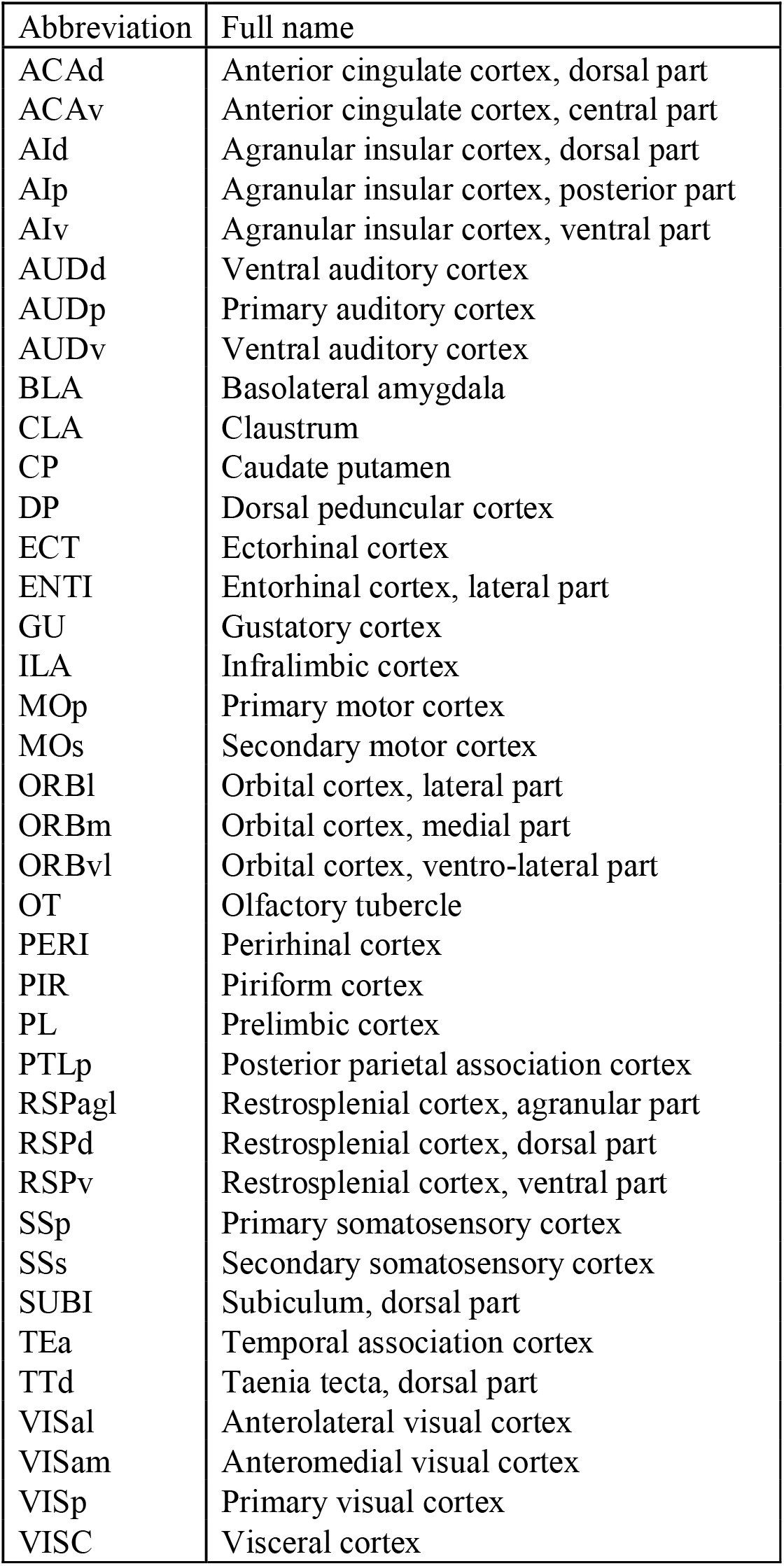
Abbreviations of brain anatomical regions used in the study.

**Supplementary table 2. Detailed statistics**

Download associated file

